# Human milk-derived extracellular vesicle treatment promotes the heat shock response in neonates with perinatal high fat diet exposure

**DOI:** 10.1101/2025.02.17.638569

**Authors:** Jasmyne A. Storm, Jueqin Lu, Mon Francis Obtial, Sanoji Wijenayake

## Abstract

Maternal consumption of a high-fat diet (mHFD) during perinatal life (the collective prenatal and postnatal periods) influences neonatal development, initiates hypothalamic-pituitary-adrenal (HPA) axis activation, and impacts the long-term physiological and metabolic health of offspring. Milk-derived extracellular vesicles (MEVs) are lipid-coated nanovesicles found in mammalian milk that survive intestinal degradation and cross complex biological barriers, including the blood-brain barrier. MEVs have known cytoprotective activity in peripheral organs; however, their pro-survival functions in response to chronic pro-inflammation stemming from early life nutrient stress remain unknown in the neonatal brain. Further, sex differences resulting from MEV treatment require investigation, as male and female neonates illicit variable responses to early life nutrient stress. We investigated whether MEVs promote the heat shock response (HSR), a principal pro-survival mechanism responsible for refolding or degrading misfolded protein aggregates through the action of heat shock protein (HSP) chaperones. We investigated the interaction between MEVs and the HSR in the liver, hypothalamus, and prefrontal cortex in male and female neonatal rats exposed to perinatal mHFD within the stress hyporesponsive period at postnatal day 11. MEV treatment robustly modulated the HSR in female neonates with the largest response recorded in the prefrontal cortex. Specifically, in the prefrontal cortex, MEV treatment led to an upregulation of the main transcription factor (HSF1), while downregulating the negative regulators of HSF1 (Hsp70 and Hsp90). These results suggest that MEVs may influence pro-survival outcomes in the prefrontal cortex by activating HSF1-mediated pro-survival in a sex specific manner in response to mHFD.

## 1.0 Background

Perinatal (the combined prenatal and postnatal periods) development is sensitive to maternal factors (Nagel et al., 2022). Maternal malnutrition resulting from over- or under-consumption of macro/micronutrients influences developmental programming and impacts the long-term health of offspring (Abuaish et al., 2020; Eisha et al., 2022; Leghi et al., 2020; Wijenayake et al., 2020). Specifically, the maternal consumption of a high saturated fat diet (mHFD) increases the risk of developing congenital defects, metabolic disorders, and neurodevelopmental and neuropsychiatric disorders in offspring (Davis and Mire, 2021; Glastras et al., 2018; Leddy et al., 2008; Makris et al., 2022; Pillai et al., 2016). Studies conducted in murine and ovine models have demonstrated that mHFD exposure during the perinatal period alters metabolism; decreases stem cell proliferation, brain weight, bone density, and muscle development; increases adiposity and bodyweight; and increases pro-inflammatory cytokine production and circulation (Bautista et al., 2016; Chen et al., 2017; Fiorotto et al., 1995; Pillai et al., 2016; Reed et al., 2014; Urbonaite et al., 2022; Wijenayake et al., 2020). These exposures may also lead to sex-specific physiological and epigenetic changes in offspring that persist into adulthood (Howie et al., 2012; Sasaki et al., 2013; Young and Ramakrishnan, 2020).

Perinatal exposure to mHFD also leads to the activation and chronic potentiation of the hypothalamic-pituitary-adrenal (HPA) stress axis (Bose et al., 2009; Chrousos, 1995; Volqvartz et al., 2023). The HPA axis is a neuroendocrine system responsible for regulating glucocorticoid levels in response to stress (Smith and Vale, 2006). Several organs are implicated in the HPA-mediated stress response in the CNS and in the periphery. The hypothalamus is the main HPA axis regulator (Sullivan and Gratton, 2002). During homeostasis, the hypothalamus regulates energy metabolism, thermoregulation, circadian cycles, and feeding behaviour (Coll and Yeo, 2013; Williams et al., 2001). During stress, however, the paraventricular nucleus (PVN) of the hypothalamus activates the HPA-mediated stress response through the release of corticotrophin releasing hormone (CRH) and arginine vasopressin (AVP) (Bao et al., 2008; Stephens and Wand, 2012). CRH and AVP stimulate the anterior pituitary gland to release adrenocorticotrophic hormone (ACTH) into general circulation and induces the release of glucocorticoids from the adrenal glands (Bao et al., 2008; Stephens and Wand, 2012). Many extrahypothalamic structures are also involved in HPA stress axis responses (Sullivan and Gratton, 2002). Lesion and glucocorticoid inhibition studies have elucidated the role of the prefrontal cortex as a site of glucocorticoid receptor-mediated feedback inhibition of the HPA axis (Diorio et al., 1993; Laryea et al., 2015; McKlveen et al., 2013; Sullivan and Gratton, 2002). Peripheral organs, including the liver, are also impacted by HPA-mediated neuroinflammation via the liver-gut-brain axis (Butterworth, 2013).

Heat shock proteins (HSPs) are molecular chaperones that are involved in the heat shock response (HSR). HSPs participate in the initiation of protein folding, refolding, repair, disaggregation, and degradation (Sottile and Nadin, 2018). In addition to their homeostatic functions, HSPs are upregulated in response to environmental stress including oxidation, heat and cold shock, and exposure to contaminants and toxins (Hoter and Naim, 2019; Ikwegbue et al., 2017; Luu et al., 2018). Interestingly, mHFD-induced metabolic dysfunction is also associated with HSR dysregulation (Habich and Sell, 2015; Hoter and Naim, 2019; Sabbah et al., 2019).

Mammalian milk is a complex, heterogeneous biological fluid tailored to meet the energetic and developmental requirements of newborns (Eisha et al., 2022; Roy et al., 2020). Milk composition is sensitive to maternal diet. Consumption of a mHFD increases pro-inflammatory cytokine levels and decrease neuroprotective molecules, such as carotenoids and long-chain polyunsaturated fatty-acids, such as docosahexaenoic acid and eicosapentaenoic acid in milk (Bautista et al., 2016; Buonfiglio et al., 2016). Despite select studies illustrating mHFD-induced changes in select milk components, maternal milk feeding is beneficial and necessary to support the healthy development of offspring, irrespective of the maternal metabolic state, as the transmission of a plethora of biologically active molecules in milk mitigate the negative developmental outcomes associated with gestational mHFD (Abuaish et al., 2020; Dai et al., 2023; Eisha et al., 2022; Leghi et al., 2020; Ozkan et al., 2020; Wijenayake et al., 2023, 2021). One type of biologically active molecule in mammalian milk is milk-derived extracellular vesicles (MEVs). MEVs are lipid-coated nanovesicles (30-150 nm in size) secreted mainly from mammary gland epithelial cells (Reif et al., 2019; Wijenayake et al., 2021) with immunomodulatory and anti-inflammatory capabilities (Lou et al., 2024). MEVs decrease rates of apoptosis, attenuate NFκB-mediated pro-inflammation, and reduce the progression of gastrointestinal diseases, including necrotizing enterocolitis and inflammatory bowel disease (Filler et al., 2023; García-Martínez et al., 2022; Jiang et al., 2023, 2021; Karra et al., 2022; Nolan et al., 2020; Tong et al., 2023; Wang et al., 2022).

Despite the known anti-inflammatory and cytoprotective potential of MEVs in peripheral organs, the ability of MEVs to interact with and enhance existing pro-survival responses require further investigation. The main objectives of our study are to 1) explore the association between MEVs and the HSR in modulating cytoprotection in offspring liver, hypothalamus, and prefrontal cortex in response to mHFD exposure, and 2) investigate these changes in male and female neonates during a critical developmental window, PND11, which coincides with the peak lactational period in rats and the stress hyporesponsive period (SHRP). The SHRP spans 14 days after birth in neonatal rodents, wherein the adrenal response to stress or adverse experience is minimal, resulting in stable levels of circulating glucocorticoids (Sapolsky and Meaney, 1986). However, mHFD has been shown to induce stress responses during the SHRP (Abuaish et al., 2018; Sasaki et al., 2014). To our knowledge, this study is the first to explore the relationship between MEVs and the HSR in male and female neonatal rats in the context of promoting cytoprotection in HPA-axis associated tissues.

## 2.0 Materials and Methods

### 2.1 MEV isolation and characterization

Unpasteurized human donor milk (n = 11 anonymous donors) was obtained from NorthernStar Mothers Milk Bank (Calgary, AB, Canada) and pooled into a homogeneous mixture to limit variability in milk composition between the donors. Serial ultracentrifugation of the milk was used to remove creams, fats, and milk fat globular membranes (twice at 3,000 x g, 10 mins, 22 °C), cellular debris (twice at 1,200 x g, 10 mins, 4 °C), and other cells (twice at 21,500 x g, 30 mins, 4 °C, then once at 21,500 x g, 60 mins, 4 °C) (Wijenayake et al., 2021). Residual debris and cells were removed by filtering the remaining whey component using a 0.45 µm syringe filter (UltiDent: 229749). Casein proteins were removed using acetic acid precipitation (1:1000, v/v), centrifugation (4,500 x g, 30 mins, 4 °C), and filtration using a 0.22 µm syringe filter (UltiDent: 229747) (Morozumi et al., 2021). MEVs were isolated from the remaining supernatant via ultracentrifugation at 100,000 x g for 120 mins at 4 °C (Beckman-Coulter XL-100; SW55T1 Swing-Bucket Rotor; no brake). The MEV pellet was washed in 1X phosphate buffered saline (PBS) (1:1, v/v), ultracentrifuged a second time at the same settings, and resuspended in filtered 1X PBS (1:2, v/v to original whey volume). The supernatant from the second ultracentrifugation was used as the EV-depleted negative control (Wijenayake et al., 2021).

MEVs were characterized in accordance with the Minimum Information for Studies of Extracellular Vesicles (MISEV) 2023 guidelines (Welsh et al., 2024). MEV particle size and concentration were quantified using nanoparticle tracking analysis (Malvern Instruments Ltd.; NanoSight NS300) at the Structural and Biophysical Core Facility at the Hospital for Sick Children (Toronto, ON, Canada). MEVs were diluted (1:300, v/v in 1x filtered PBS) and captured using an sCMOS camera (level 15, detection threshold 10, 532nm green laser, 30s capture speeds, 3x replicates) as per Wijenayake et al. (2021). MEV morphology and integrity was characterized using transmission electron microscopy (FEI Talos F200x S/TEM) at the Manitoba Institute of Materials (Winnipeg, MB, Canada). MEVs were suspended in 1x PBS (1:3, v/v), loaded onto 400 mesh carbon-coated formvar film copper grids (Electron Microscopy Sciences: CF400-CU-50), and negatively stained using 2% uranyl acetate as per Wijenayake et al. (2021). MEV biomarkers were characterized using western immunoblotting, where the presence of three endosome-specific proteins (tetraspanins, CD9 and CD81, and an EV-biogenesis factor, syntenin-1) were detected. Protein abundance of a cellular marker of the endoplasmic reticulum (calnexin) was used as negative control and proteins isolated from human microglia cell lineage 3 (HMC3) cells were used as a cellular control. The results of the MEV characterization are available in Storm et al. (2025). The same batch of MEVs isolated from the same 11 milk donors were used across studies.

### 2.2 Animal care and treatment

All experimental procedures were approved by the University of Winnipeg Animal Care Committee (AE18072) and were in accordance with the guidelines of the Canadian Council on Animal Care. 7-week-old female (n = 12) and male Long Evans rats (n = 6) (strain code: 006) were purchased from Charles River Canada (St. Constant, QC). All rats were housed in same-sex pairs until mating and maintained on a 12h:12h light/dark cycle with *ad libitum* access to food and water. Following one-week of acclimatization, a subset of females (n = 6) was placed on a mCHD consisting of 10% kcal fat (Research Diet Inc., D12450J), with the remaining females (n = 6) placed on a mHFD consisting of 60% kcal fat (Research Diet Inc., D12492) (**Figure 1**). The mCHD was matched in sucrose content to the mHFD. Diet was maintained for 4 weeks prior to mating, and throughout gestation and lactation (Abuaish et al., 2020; Sasaki et al., 2014, 2013; Wijenayake et al., 2020). For breeding, females were pair housed with one male for seven days, after which females were housed individually throughout gestation and lactation. Pregnancy was inferred by monitoring daily body weight gain rather than using vaginal smears, allowing for reduced manipulation and maternal stress for dams. Vaginal plugs were not used because they do not guarantee pregnancy (Abuaish et al., 2018). Dam weight gain and caloric intake was measured once a week prior to mating, and throughout gestation, then daily during lactation. Day of parturition is considered PND0 and the day before is considered the last day of gestation. At PND2 litters were weighed and culled to 12 neonates/litter (n = 6 females and n = 6 males, where possible) to standardize maternal care provisions across litters. Maternal care differences resulting from consuming this specific HFD is reported to be negligible (Abuaish et al., 2018; Rahman et al., 2022).

**Figure 1.**
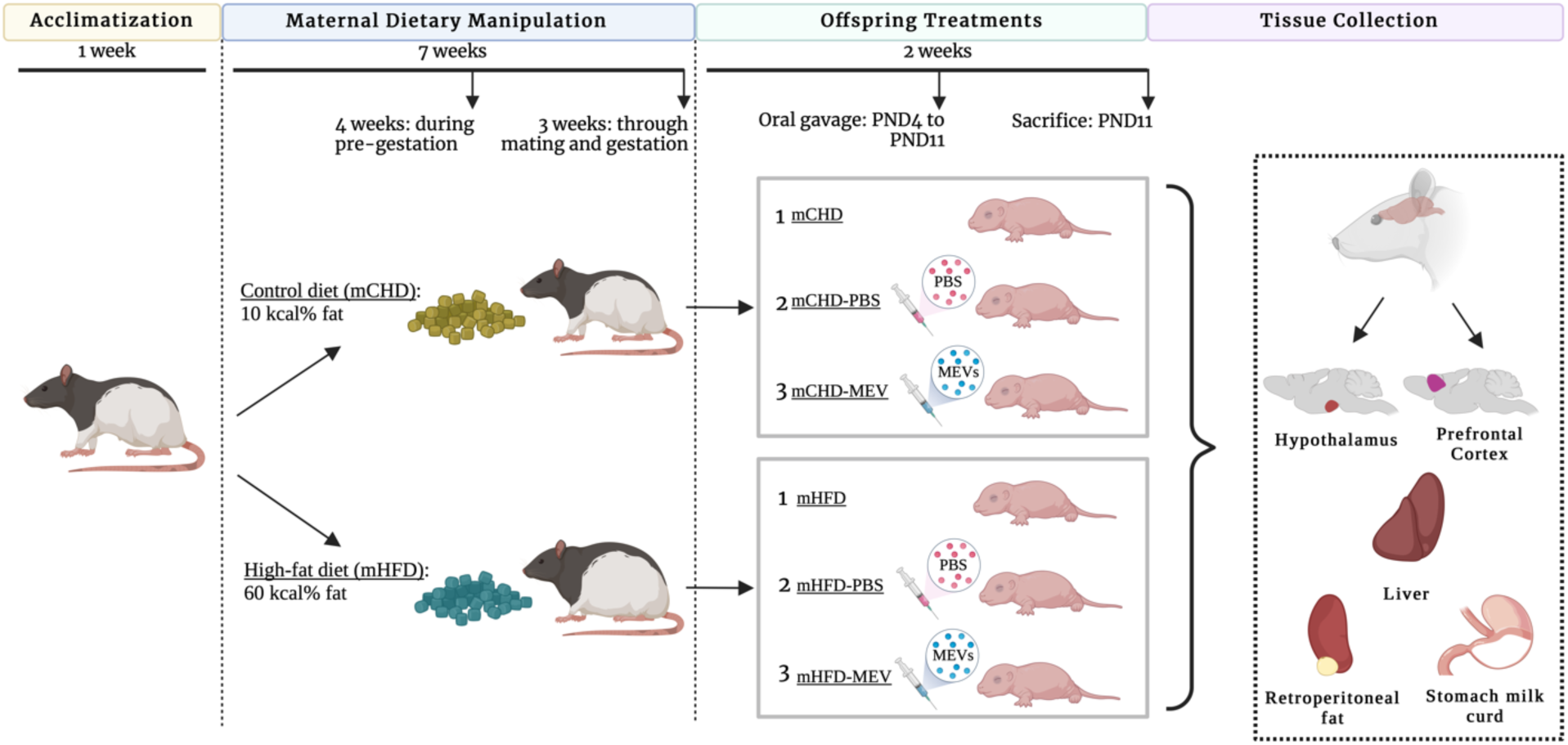
Animal care and study overview. Following one week of acclimatization, dams were placed on a control diet (mCHD) consisting of 10% kcal fat, or a high-fat diet (mHFD) consisting of 60% kcal fat, (n = 6/diet). Diet was maintained for four weeks prior to mating, throughout mating and gestation, and lactation. After parturition, at postnatal day (PND) 2, litters were weighed and culled to 12 pups/litter (n = 6 females and n = 6 males, where possible) to standardize maternal care provisions across litters. MEV treatment began at PND4, where a subset of neonates were controls (mCHD or mHFD) or received a vehicle gavage (mCHD-PBS or mHFD-PBS) or received MEV gavage (mCHD-MEV or mHFD-MEV) (n = 1-2 pups/litter/sex). Oral gavage (1×10^10^ MEVs/g of body weight) was administered twice a day, 6h apart, until PND11. At sacrifice on PND11, liver, hypothalamus, prefrontal cortex, stomach milk curd, and retroperitoneal fat were collected for downstream molecular analysis.

MEV treatment via oral gavage began at PND4 and continued until the end of the study at PND11. MEVs were administered using disposable 22-gauge polypropylene feeding tubes (Instech Laboratories: FTP-22-25). n = 1-2 neonates/litter/sex were randomly assigned to three treatment groups, depending on sex ratio and litter sizes: 1) neonates that were separated from the nest with their littermates and handled but did not receive gavage (referred to as mCHD or mHFD), 2) neonates that received the vehicle control of 1X PBS (referred to as mCHD-PBS or mHFD-PBS), and 3) neonates that received 1×10^10^ MEVs in 1X PBS per gram of bodyweight (referred to as mCHD-MEV or mHFD-MEV). Oral gavage was administered twice a day, during the light phase, 6h apart. The oral gavage dosage used in our study is similar to previous studies (Hock et al., 2017; Li et al., 2019; Manca et al., 2018). MEVs have a high cross-species tolerance and the administration of human and/or bovine MEVs to murine species does not affect viability or induce physiological effects (Manca et al., 2018; Mondal et al., 2023; Zhong et al., 2021). Due to the large volumes of rodent milk that is required to isolate sufficient MEVs to gavage neonates twice a day, for 7 days, it was not feasible to use Long Evans rat milk for this experiment. The neonates were separated from the nest for a maximum of 15 minutes/procedure and 30 minutes/day. As per previously published works, an elevation in plasma corticosterone (ng/ml) was not observed until >2h maternal separation (Kuhn et al., 1990; Levine et al., 1991; Tiba et al., 2004). Pup body weight and naso-anal length were determined daily from PND2 to PND11. The Lee index is used as an index of obesity in rodents and has been shown to correlate to carcass fat percentage (Simson and Gold, 1982). The Lee index was calculated as follows as per Simson and Gold (1982):

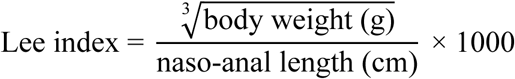

At PND11 neonates were euthanized via swift decapitation. The liver, hypothalamus, prefrontal cortex, stomach milk curd, and retroperitoneal fat were collected, flash frozen, and stored at −80 °C. In total, 48 female and male neonates at PND 11 from n = 8 separate litters were used for this study. n = 2 females and n = 2 males were used per litter.

### 2.3 RNA extraction and cDNA synthesis

Total soluble RNA (≥ 18 nucleotides) was extracted from the liver, hypothalamus, and prefrontal cortex (n = 4 biological replicates/diet/treatment/sex) using TRIzol reagent (ThermoFisher Scientific: 15596018) according to the manufacturer’s instructions. RNA samples were resolved on a 1% TAE-agarose gel with 2x RNA loading dye (1:1, v/v) (Life Technologies: R0641) stained with Red Safe dye (FroggaBio: 21141) to verify stability and integrity (**Figure S1-S3**). RNA concentration (ng/µL) and purity (A260:280 and A260:230 ratios) were determined using a Nanodrop One/One^C^ Microvolume-UV/Vis spectrophotometer (ThermoFisher Scientific: ND-ONE-W). Genomic DNA was removed using DNase I treatment, and an RNA Clean & Concentrator kit (Zymo Research: R1017) was used to clean select RNA samples, according to the manufacturer’s instructions. Samples with an A260:280 ratio of 1.8-2.0 were used for cDNA synthesis. 2000 ng of total soluble RNA was reverse transcribed into cDNA using the High-Capacity cDNA Reverse Transcription kit (Applied Biosystems: 4368814) according to the manufacturer’s instructions. cDNA was synthesized using a T100™ Thermal Cycler (Bio-Rad: 1861096) (amplification parameters: 25 °C for 10 mins; 37 °C for 120 mins; 85 °C for 5 mins; hold at 4 °C). Samples were stored at −80 °C for future use.

### 2.4 Primer design and RT-qPCR

Gene expression of HSF1, *HSPA1B*, *HSP90AA1*, and *DNAJB1* were measured via RT-qPCR using a QuantStudio™ 5 PCR system (Applied Biosystems: 96-well and 0.2 mL block) using Fast SYBR™ Green Master Mix chemistry (Applied Biosystems: 4385612). Internal controls were *GAPDH* and *YWAZ* (liver), *GUSB* (hypothalamus), and *18S* rRNA and *YWAZ* (prefrontal cortex). Additional candidate internal controls were tested in the liver (*18S rRNA*, *Actin*-*β*), hypothalamus (*18S rRNA*, *Actin*-*β, B2M, Eif1a1, GAPDH, GDI1, Nupl2, PGC1α, PPIA, Rpl5, Rpl27, Rpl32, SDHA, TBP, YWAZ*), and prefrontal cortex (*Actin*-*β, GAPDH*), but were determined unsuitable for normalization because they varied across treatment or diet in either sex (**Table S1)**. mRNA-specific primers were designed using nucleotide sequence information from NCBI or obtained from previous literature. Sequence information from NCBI and the OligoAnalyzer™ Tool from Integrated DNA Technologies (IDT, Coralville, Iowa, USA) were used to assess primer compliance with the Minimum Information for Publication of Quantitative Real-time PCR Experiments (MIQE) guidelines (Bustin et al., 2009). Primer parameters are listed in **Table S2,** and primer bioinformatics are listed in **Table S3**.

Annealing temperatures of primer pairs were determined by testing a range of temperatures ± 5 °C of the forward primer (5’-3’) melting temperature using the QuantStudio™ 5 PCR system Veriflex settings. Temperature testing was conducted using a pool sample (10 ng/µL) representing all test samples across tissues. Melt curve analysis was conducted to determine primer pair specificity. Primer pairs that generated a single, sharp peak with a derivative reporter >200,000 and devoid of primer dimers were used for quantification. Optimal cDNA loading amounts were determined per primer pair and per tissue using an 8-point standard curve ranging from 500 ng/µL to 3.91 ng/µL. Analyses were conducted in triplicate and an inter-plate converter was used across plates to account for variability between plates and amplifications.

### 2.5 Protein extraction and analysis of abundance by western immunoblotting

Total soluble protein (n = 4 biological replicates/diet/treatment/sex) was extracted using 1x Milliplex lysis buffer (1:4 w/v of tissue sample) (Millipore-Sigma: 43-040) with 1 mM sodium orthovanadate (BioShop: SOV850.25), 10 mM sodium fluoride (BioShop: SFL001), 10 mM β-glycerophosphate disodium pentahydrate (BioShop: GYP001.50), and protease inhibitor cocktail (BioShop: PIC001.1) according to Tessier et al. (2017). Animals used in the protein analysis are littermates of the animals that were used in the RNA analysis. Samples were manually homogenized using a dounce pestle in lysis buffer, then incubated for 30 minutes on ice with intermittent vortexing, followed by centrifugation (14,000 x g, 20 min, 4 °C). Protein concentration was determined using a BCA assay (Pierce™ BCA Protein assay; ThermoFisher Scientific: 23227), according to the manufacturer’s instructions, and absorbance readings were obtained using a BioTek Synergy H1 Multimode microplate reader (Agilent: BTSH1M2SI) at 562 nm. Lysates were combined with bromophenol blue loading dye with SDS (100 mM Tris-base; 4% w/v, SDS; 20% v/v, glycerol; 0.2% w/v, bromophenol blue) (Bioshop: SDS001.1) and β-mercaptoethanol (10%, v/v) (Bioshop: MERC002.500), then vortexed and heated to 95 °C for 10 minutes for denaturation. Samples were stored at −20 °C for future use. The same methodology and parameters used for protein extraction from rat liver, hypothalamus and prefrontal cortex were also used to isolate soluble protein from MEV pellets for characterization.

Protein lysates were resolved using 10-15% SDS-polyacrylamide gels. A standard curve (5μg - 35 μg) using an organ-specific pool sample was used to determine optimal protein amount to load per antibody and per tissue. PageRuler™ Plus prestained protein ladder (ThermoFisher Scientific: 26619) was used as a molecular weight standard, and a liver pool was run in duplicates as the interblot converter to be used for normalization and comparisons across immunoblots. Samples were resolved (n = 3-4/treatment group/sex/diet) on SDS-polyacrylamide gels for 110 minutes in 1x Tris-Glycine running buffer (0.3%, w/v Tris-base; 14.4%, w/v glycine; 1%, w/v SDS) at 180 V in a Sub-Cell GT electrophoresis cell (Bio-Rad: 1704401). Proteins were transferred onto 0.45 µm PVDF membranes (Bio-Rad: 1620174) using a Trans-Blot Turbo Transfer System (Bio-Rad: 1704150). Membranes were blocked using 1-20% casein-TBST solution (30 minutes, 22 °C), incubated with primary antibody (1:500 or 1:1000, v/v), followed by goat HRP-conjugated anti-rabbit IgG secondary antibody (1:10,000 or 1:15,000, v/v) for 45 minutes at 22 °C, with all steps performed on a rocker. The immunoblots were visualized using western ECL substrate solution (ThermoFisher Scientific: 34580) and chemiluminescence imaging (ChemiDoc™ MP imaging system; Bio-Rad: 12003154) with Image Lab Software (version 6.1). The immunoblots were stained for 2 minutes in Coomassie brilliant blue (BioShop: CBB555.10) solution (0.25%, w/v Coomassie blue salt; 7.5% acetic acid; 50%, v/v methanol) and destained for 5 minutes with destain solution (25%, v/v methanol; 10%, v/v acetic acid) at room temperature, on a rocker. ImageJ (version 1.5.3) was used to quantify protein abundance as per Abràmoff et al. (2004). Western immunoblotting parameters for HSR targets are listed in **Table S4** and relevant antibody information in **Table S5**. ECL and Coomassie images for HSR targets are displayed in **Figure S4-S7**.

### 2.6 Statistical analysis

Statistical analysis was carried out using SPSS version 29.0.2.0 (IBM Corp.) and figures were constructed using GraphPad Prism 7 and BioRender.

Normality was assessed using Shapiro-Wilk tests for datasets with n < 30. Extreme outliers with an IQR > 3 were identified using the SPSS boxplot outlier function and removed from the dataset when data was not normally distributed. Data that did not achieve normality was analyzed with Mann Whitney U (for 2 comparisons) and Kruskal Wallis H tests followed by Dunn’s post hoc tests (for 3 or more comparisons).

Factorial (time, diet [mCHD or mHFD], treatment [control, PBS, MEV], sex) repeated-measures general linear model (GLM) was used with Bonferroni correction for maternal bodyweight and average caloric intake during pre-gestation, gestation, and lactation, and offspring bodyweight and Lee-index. Greenhouse-Geisser corrections were used as the datasets failed Mauchly’s tests of sphericity. Dam mating success, litter size, and postnatal weight gain were determined using two-tailed, independent samples t-tests to compare mCHD and mHFD litters.

Univariate GLM (diet, treatment, sex) was conducted to analyze total retroperitoneal fat weight and stomach curd weight, as well as for parametric RT-qPCR and western immunoblotting data in offspring. Tukey HSD post hoc test was used to conduct pairwise comparisons between treatment groups. A Pearson correlation was used to compare offspring bodyweight (g) and retroperitoneal fat weight (g), and offspring bodyweight and stomach milk curd weight (g).

All RT-qPCR and western immunoblotting data were analyzed with sexes separated, as perinatal mHFD exposure leads to sex differences (Abuaish et al., 2018; Sasaki et al., 2013; Wijenayake et al., 2020). n = 2 neonates per litter per sex were used for transcript and protein analyses, as such dam ID was not used as a covariate. The core results of the study compare male and female offspring across two diets (mCHD and mHFD) and two treatments (controls and MEV gavaged). The dataset for neonates who received the 1X PBS vehicle is in **Table S6**. A p-value < 0.05 was considered statistically significant for all statistical tests. Data are represented as mean ± SEM.

## 3.0 Results

### 3.1 Changes in dam bodyweight and caloric intake from pre-gestation and gestation throughout lactation, and analysis of mating success, litter size, and postnatal weight change

Dam bodyweight remained unchanged throughout pre-gestation, gestation, and lactation between mCHD and mHFD (**Figure 2a**). Specifically, there were no significant differences in bodyweight between mCHD and mHFD dams (*F*_(1,9)_ = 0.230, p = 0.643), nor significant interactions between the diet conditions and weeks spent on respective diet regimes (*F*_(1.882, 16.941)_ = 0.357, p = 0.693). Dam body weight increased with age in mCHD and mHFD (*F*_(1.882, 16.941)_ = 82.507, p < 0.001). Similarly, there were no significant changes between mCHD and mHFD dams in mating success (66.7% success for mCHD and mHFD), sizes of litters (*t*(8) = −1.452, p = 0.185), or postnatal weight change (*t*(8) = −-1.549, p = 0.160) (**Figure 2b-d**). Average daily caloric intake of dams was analyzed throughout pre-gestation, gestation, and lactation (**Figure 2e**). mHFD dams had a higher average daily caloric intake than mCHD dams (*F*_(1,9)_ = 13.079, p = 0.006), where mHFD dams consumed more kCal per day in the first and second weeks of pre-gestation. Caloric intake also increased over time in both mCHD and mHFD dams (*F*_(1.313, 11.820)_ = 32.726, p < 0.001), although there was no significant interaction between diet conditions and weeks spent on the respective diets (*F*_(1.313, 11.820)_ = 2.075, p = 0.175). Daily caloric intake during the lactational period was also measured in dams post-parturition (**Figure 2f**). During lactation, there are no significant differences in daily caloric intake between mCHD and mHFD dams (*F*_(1, 8)_ = 2.674, p = 0.141); however, as seen previously, there is an increase in caloric intake over time (*F*_(1.921, 15.364)_ = 3.835, p = 0.046). No significant interactions between diet conditions and days spent on the diet during lactation were observed (*F*_(1.921, 15.364)_ = 1.749, p = 0.207).

**Figure 2.**
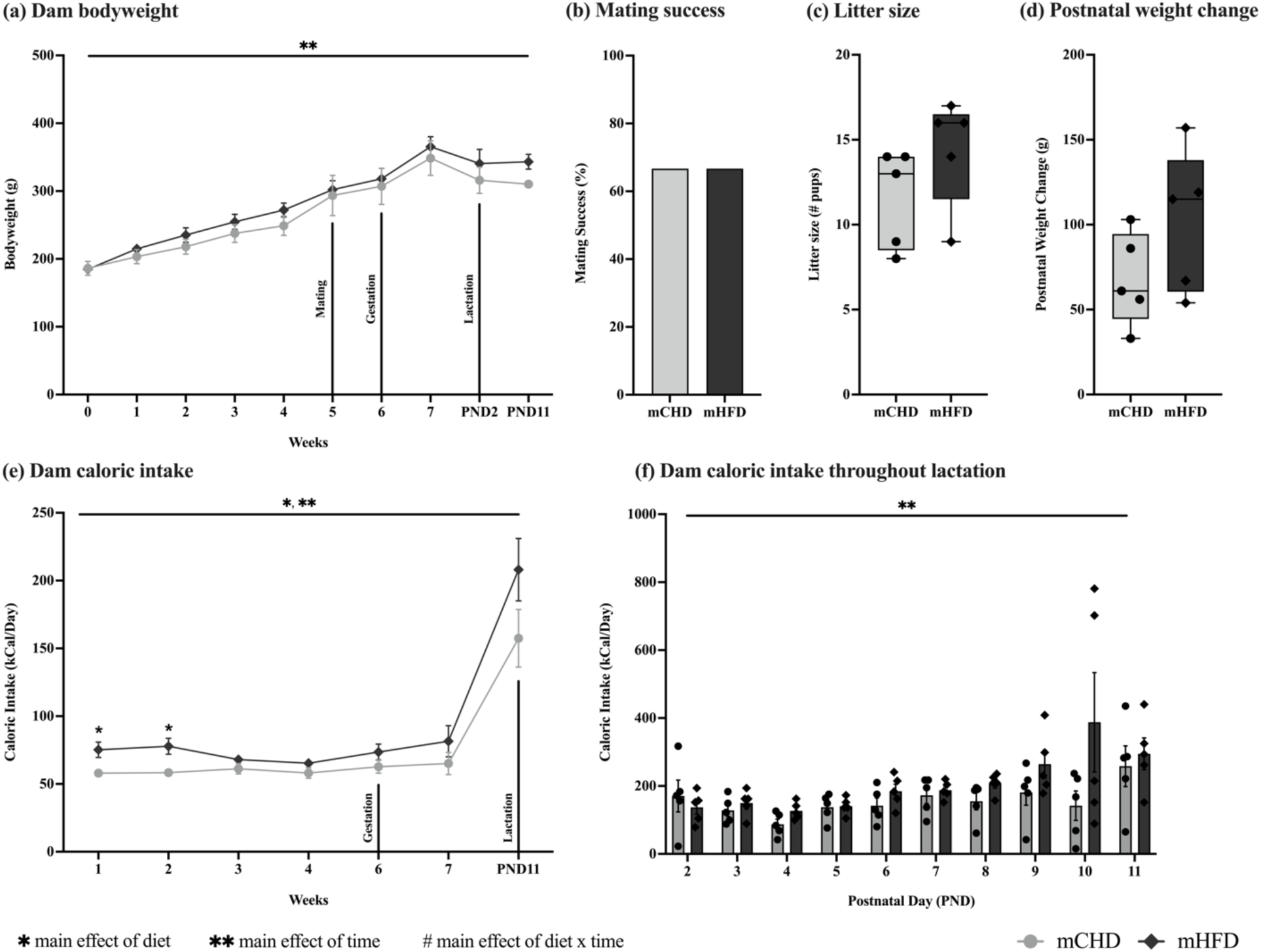
Maternal responses to high-fat diet (mHFD) consumption. **(a)** Changes in dam bodyweight during pre-gestation, gestation, and lactation, between control diet (mCHD) and mHFD (n = 6/diet). **(b)** Mating success of mCHD and mHFD dams. **(c)** Litter sizes at parturition. **(d)** Dam postnatal weight change between parturition and the end of the study at postnatal day (PND) 11. **(e)** Average daily caloric intake (kCal/day) in dams during pre-gestation, gestation, and lactation. **(f)** Daily caloric intake (kCal/day) in dams throughout lactation. *Main effect of diet (p<0.05), **Main effect of time (p<0.05), and ^#^Interaction between diet x time (p<0.05). Data presented are means ± standard error.

### 3.2 Change in offspring bodyweight and Lee index from PND2 to PND11

Changes in offspring bodyweight were analyzed from PND2 until sacrifice at PND11 (**Figure 3a**). No differences in bodyweight were observed between male and female neonates within a diet (*F*_(1, 98)_ = 2.835, p = 0.095) or between treatment groups (*F*_(2, 98)_ = 0.240, p = 0.787); therefore, changes in offspring bodyweight were determined without separating neonates by sex or treatment. Exposure to mHFD influenced offspring weight and caloric intake. Offspring bodyweight increased throughout the lactational period (*F*_(1.369, 142.428)_ = 2086.469, p < 0.001) with significantly higher bodyweight in neonates borne to mHFD dams compared to mCHD dams (*F*_(1, 104)_ = 9.962, p = 0.002). Furthermore, a significant interaction between days and diet was observed (*F*_(1.369, 142.428)_ = 35.396, p < 0.001). No significant interactions were observed between days and treatment group (*F*_(2.739, 142.428)_ = 0.872, p = 0.449), nor days with diet and treatment (*F*_(2.739, 142.428)_ = 1.210, p = 0.307).

**Figure 3.**
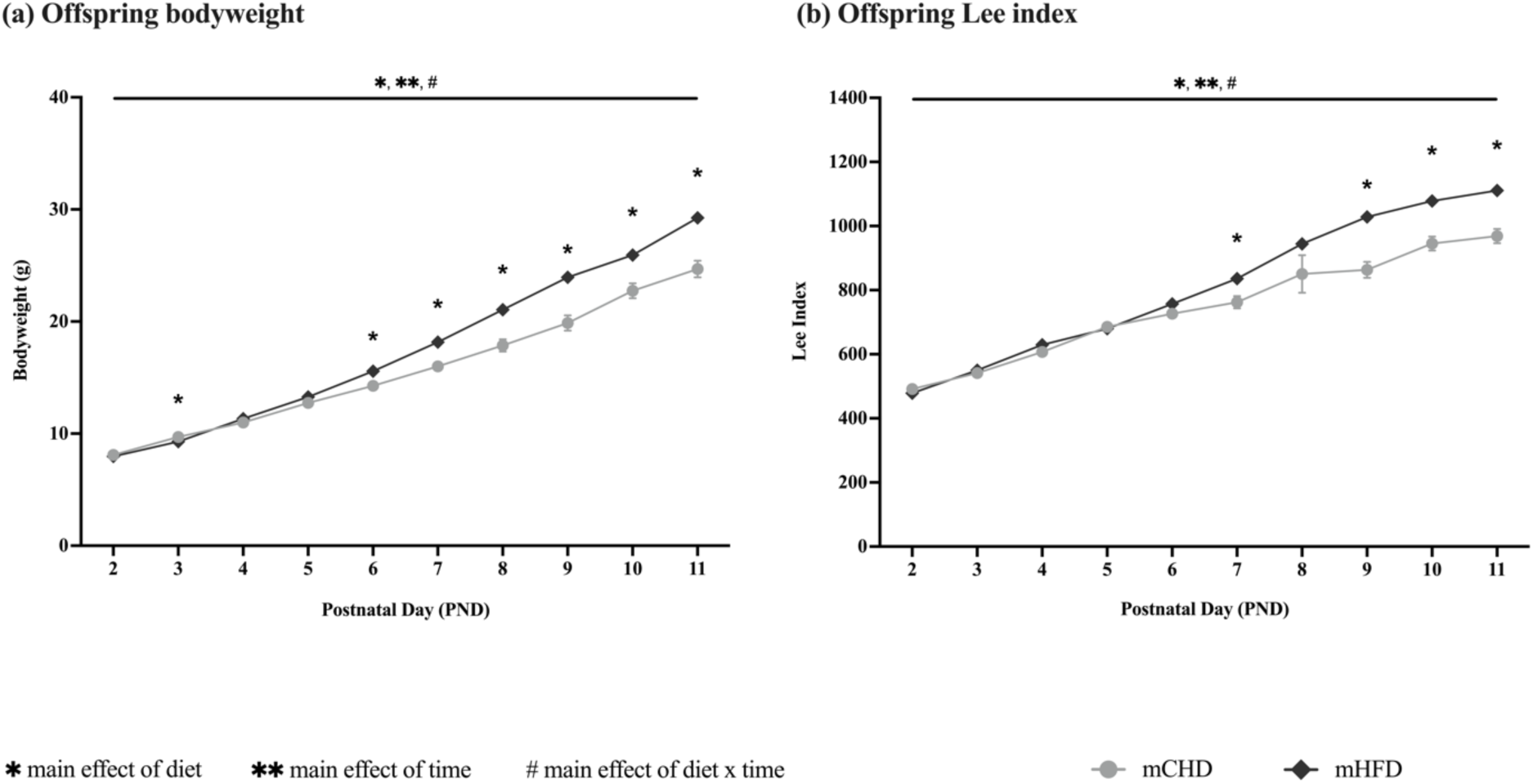
Offspring responses to maternal diet. **(a)** Changes in offspring bodyweight from postnatal day (PND) 2 to the end of the study at PND11 in response to control diet (mCHD) and mHFD (n = 24-31/diet/sex). **(b)** Changes in offspring Lee index throughout lactation between pups in mCHD and mHFD diet groups. *Main effect of diet (p<0.05), **Main effect of time (p<0.05), and ^#^Interaction between diet x time (p<0.05). Data were combined across sex and treatment as no main effects were seen. Data presented are means ± standard error.

Similar to bodyweight, changes in offspring Lee index were also analyzed from PND2 to PND11 (**Figure 3b**). No differences in Lee index were observed between male and female neonates within a diet (*F*_(1, 98)_ = 2.423, p = 0.123) or between treatment groups (*F*_(2, 98)_ = 0.163, p = 0.850); therefore, changes in offspring Lee index were also determined without separating neonates by sex or treatment. Offspring Lee index increased throughout the lactational period (*F*_(1.758, 182.823)_ = 383.968, p < 0.001) with significantly higher Lee index in neonates borne to mHFD dams than mCHD dams (*F*_(1, 104)_ = 8.357, p = 0.005). Additionally, a significant interaction between days and diet was observed (*F*_(1.758, 182.823)_ = 9.885, p < 0.001), although there were no significant interactions between days and treatment (*F*_(3.516, 182.823)_ = 0.937, p = 0.435), nor days with diet and treatment (*F*_(3.516, 182.823)_ = 1.082, p = 0.363).

### 3.3 Correlations between pup bodyweight, stomach milk curd weight and retroperitoneal fat weight

At PND11, mHFD offspring had significantly higher total retroperitoneal fat weight than mCHD offspring (*U* = 1873.500, p = 0.002), however, there were no significant differences in fat deposition between males and females within a diet (*U* = 1599.500, p = 0.217) or between treatment groups (*H*(2) = 0.035, p = 0.982) (**Figure 4a**). Therefore, changes were determined without separating neonates by sex or treatment. A positive linear correlation was observed between increased bodyweight and increased retroperitoneal fat weight at PND11 (R^2^ = 0.3085, p < 0.0001) (**Figure 4b**).

**Figure 4.**
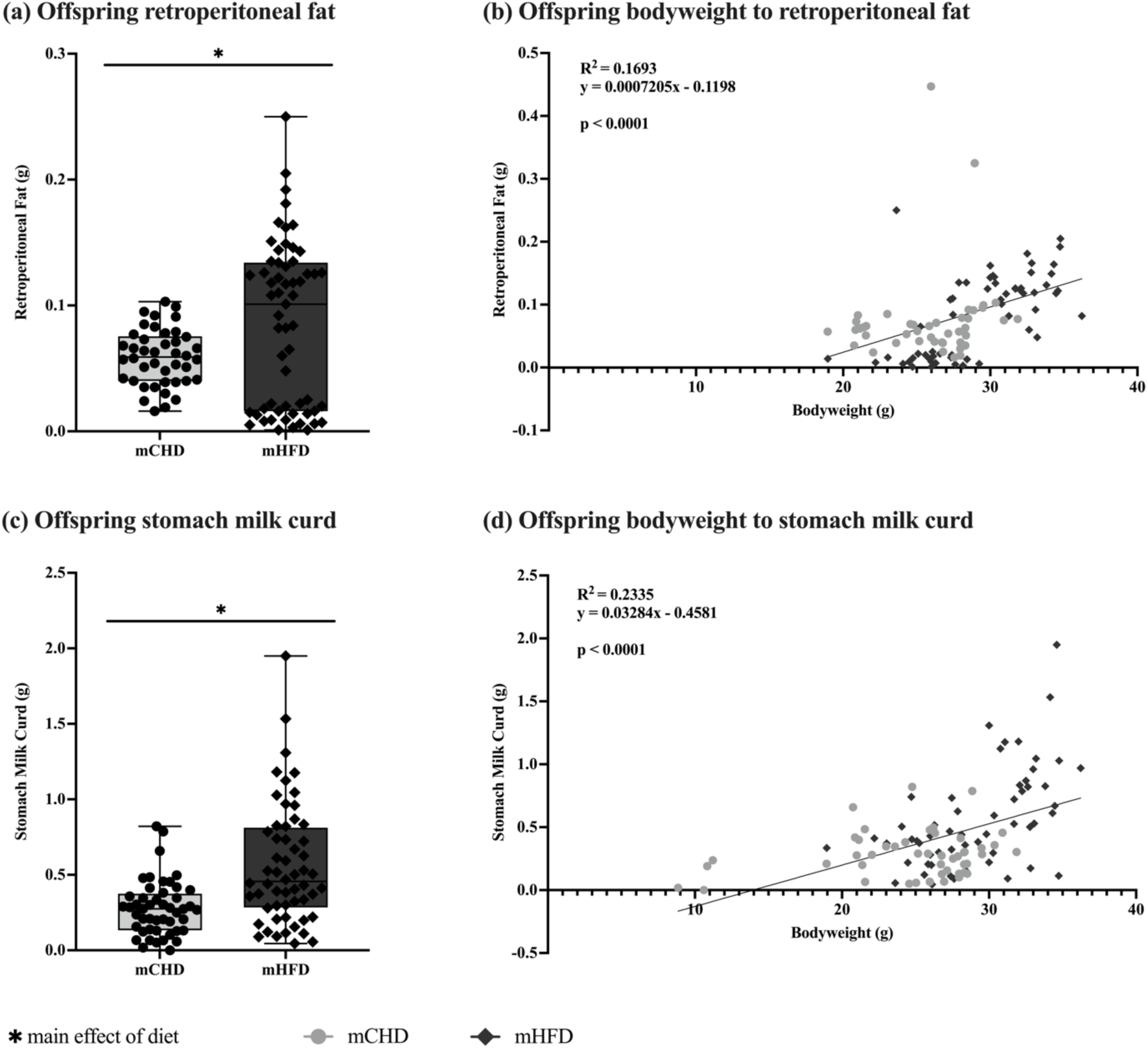
Analysis of offspring retroperitoneal fat and stomach milk curd weight in response to maternal diet. **(a)** Total retroperitoneal fat weight in neonates at postnatal day (PND) 11 in response to maternal control diet (mCHD) and maternal high fat diet (mHFD). **(b)** Pearson correlation between offspring bodyweight and amount of retroperitoneal fat. **(c)** Total stomach milk curd weight in neonates at PND11 in response to mCHD and mHFD. **(d)** Pearson correlation between offspring bodyweight and amount of stomach milk curd present in stomach. *Main effect of diet (p<0.05), **Main effect of time (p<0.05), and ^#^Interaction between diet x time (p<0.05). Data were combined across sex and treatment as no main effects were seen. Data presented are means ± standard error.

mHFD offspring also had significantly higher stomach milk curd weight at PND11 when compared to mCHD offspring (*U* = 2069.000, p < 0.001), although there were no significant differences in stomach milk weight between males and females (*U* = 1532.000, p = 0.776) or between treatment groups (*H*(2) = 3.604, p = 0.165) (**Figure 4c**). Therefore, changes were determined without separating neonates by sex or treatment. A positive linear correlation was also observed between increased bodyweight and increased stomach milk curd weight at PND11 (R^2^ = 0.2221, p < 0.0001) (**Figure 4d**).

### 3.4 Transcript abundance of HSR targets in the liver

In the liver, transcript abundance of HSF1 remained unchanged in male neonates (diet: (*F*_(1,11)_ = 1.993, p = 0.186); treatment: (*F*_(3,11)_ = 1.713, p = 0.222)) (**Figure 5a**). In female neonates, while no diet effect was observed in HSF1 transcript (*F*_(1,12)_ = 0.007, p = 0.933), there was an effect of MEV treatment (*F*_(3,12)_ = 7.683, p = 0.004) with higher HSF1 transcript in mCHD-MEV neonates compared to mCHD (Tukey HSD, p = 0.006), lower HSF1 transcript in mHFD-MEV compared to mCHD-MEV (Tukey HSD, p = 0.034), and higher HSF1 transcript in mHFD compared to mCHD (Tukey HSD, p = 0.027) (**Figure 5a**).

**Figure 5.**
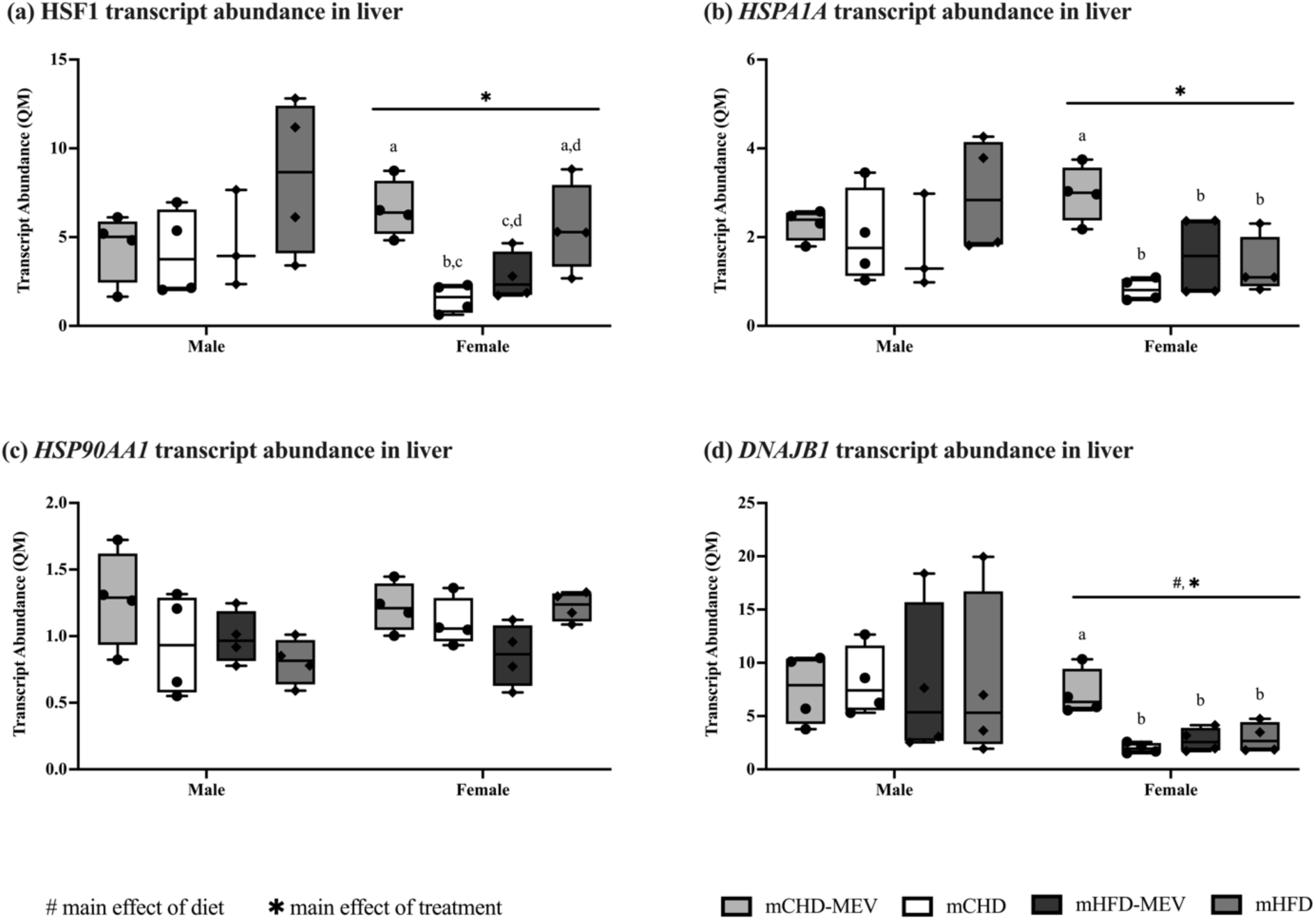
Transcript abundance of the heat shock response genes in the liver of male and female neonates at postnatal day (PND) 11 as determined by RT-qPCR. **(a)** HSF1 liver transcript abundance. **(b)** *HSPA1A* liver transcript abundance. **(c)** *HSP90AA1* liver transcript. **(d)** *DNAJB1* liver transcript abundance. Quantity means are normalized to the geometric mean of two reference genes with stable expression: *GAPDH* and *YWAZ*. ^#^Main effect of diet (p<0.05). *Main effect of MEV treatment (p<0.05). Pairwise comparisons between treatment groups are indicated with lowercase letters, where significant differences (p<0.05) are denoted by different letters. mCHD: Neonates born to mCHD dams that did not receive MEV supplementation. mCHD-MEV: Neonates born to mCHD dams that received MEV supplementation. mHFD: Neonates born to mHFD dams that did not receive MEV supplementation. mHFD-MEV: Neonates born to mHFD dams that received MEV supplementation. n = 3-4 biological replicates/diet/treatment/sex. Data presented are means ± standard error.

*HSPA1A* transcript abundance remained unchanged in male neonates (diet: (*F*_(1,11)_ = 0.146, p = 0.710); treatment: (*F*_(3,11)_ = 0.961, p = 0.445)) (**Figure 5b**). In female neonates, there was no effect of diet (*F*_(1,12)_ = 1.863, p = 0.197); however there was an effect of treatment (*F*_(3,12)_ = 7.808, p = 0.004) with higher *HSPA1A* transcript in mCHD-MEV neonates compared to mCHD (Tukey HSD, p = 0.003), lower *HSPA1A* transcript in mHFD-MEV compared to mCHD-MEV (Tukey HSD, p = 0.047) and lower *HSPA1A* transcript in mHFD compared to mCHD-MEV (Tukey HSD, p = 0.019) (**Figure 5b**).

*HSP90AA1* transcript abundance remained unchanged for male neonates (diet: (*F*_(1,12)_ = 1.966, p = 0.186); treatment: (*F*_(3,12)_ = 1.819, p = 0.197)) (**Figure 5c**). In female neonates, there was no effect of diet (*F*_(1,12)_ = 1.696, p = 0.217) on *HSP90AA1* transcript. A treatment effect was observed (*F*_(3,12)_ = 3.478, p = 0.050), although individual differences between treatments were not observed (**Figure 5c**).

*DNAJB1* transcript abundance remained unchanged in male neonates (diet: (*F*_(1,12)_ = 0.003, p = 0.956); treatment: (*F*_(3,12)_ = 0.011, p = 0.998)) (**Figure 5d**). In female neonates, *DNAJB1* transcript was responsive to changes in diet (*F*_(1,11)_ = 4.917, p = 0.049). There was also a treatment effect (*F*_(3,11)_ = 10.080, p = 0.002), with higher *DNAJB1* transcript in mCHD-MEV compared to mCHD (Tukey HSD, p = 0.001), mHFD (Tukey HSD, p = 0.010) and mHFD-MEV (Tukey HSD, p = 0.006), respectively (**Figure 5d**). Transcript data of male and female neonates that received PBS gavage (the sham controls) is included in **Table S6**.

### 3.5 Protein abundance of HSR targets in the liver

In the liver, HSF1 protein abundance in male neonates remained unchanged (diet: (*F*_(1,12)_ = 0.499, p = 0.494); treatment: (*F*_(3,12)_ = 2.538, p = 0.106)) (**Figure 6a**). HSF1 protein abundance also remained unchanged in female neonates (diet: (*F*_(1,12)_ = 2.597, p = 0.133); treatment: (*F*_(3,12)_ = 2.965, p = 0.075)) (**Figure 6a**).

**Figure 6.**
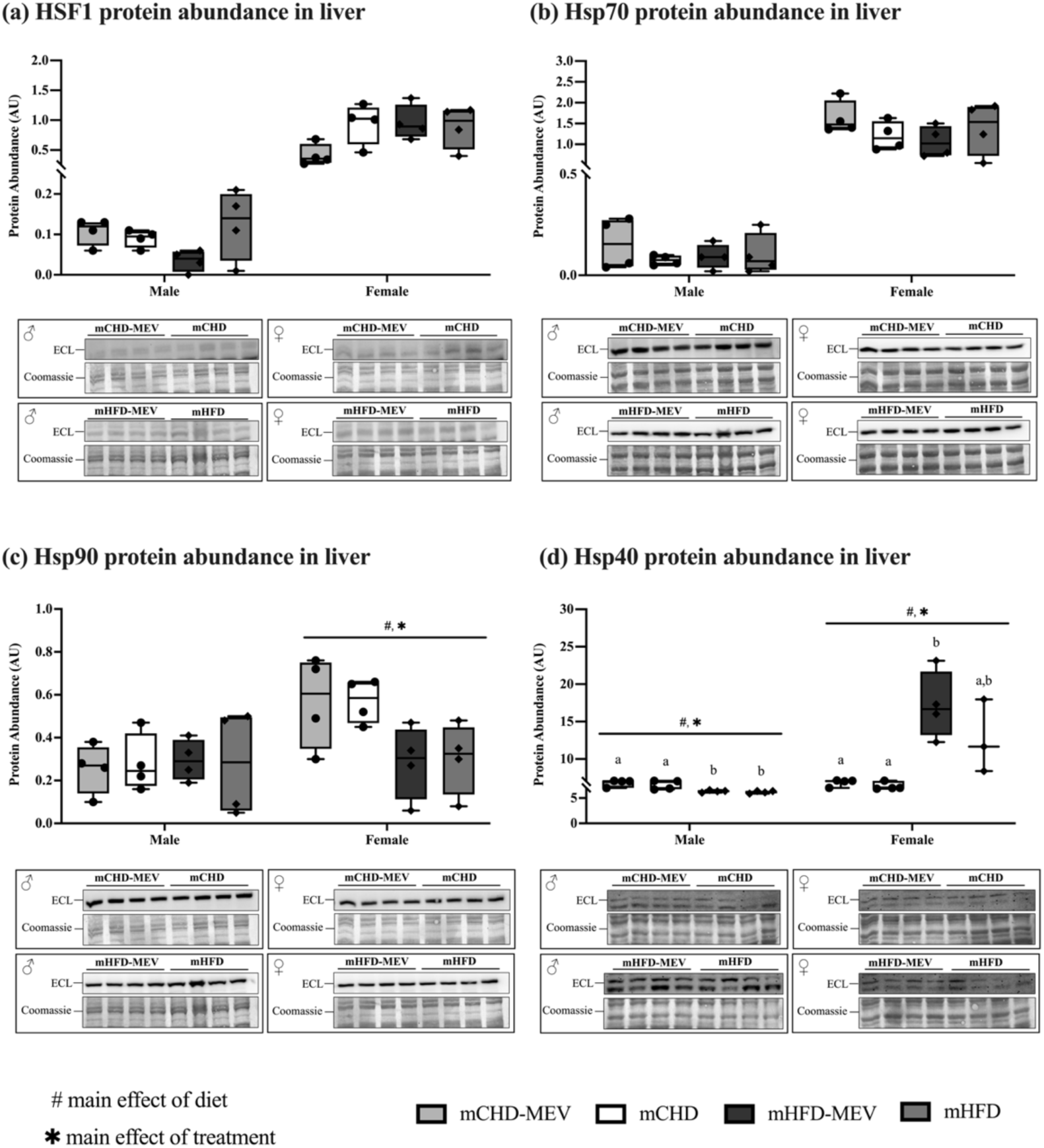
Protein abundance of the heat shock response targets in the liver of male and female neonates at postnatal day (PND)11 as determined by western immunoblotting. (a) HSF1 liver protein abundance. (b) Hsp70 liver protein abundance. (c) Hsp90 liver protein abundance. (d) Hsp40 liver protein abundance. Protein targets are normalized to the abundance of total soluble proteins in the samples using Coomassie staining. ECL and Coomassie-stained images are displayed. ^#^Main effect of diet (p<0.05). *Main effect of treatment (p<0.05). Pairwise comparisons between treatment groups are indicated with lowercase letters, where significant differences (p<0.05) are denoted by different letters. mCHD: Neonates born to mCHD dams that did not receive MEV supplementation. mCHD-MEV: Neonates born to mCHD dams that received MEV supplementation. mHFD: Neonates born to mHFD dams that did not receive MEV supplementation. mHFD-MEV: Neonates born to mHFD dams that received MEV supplementation. n = 3-4 biological replicates/diet/treatment/sex. Data presented are means ± standard error.

Hsp70 protein abundance remained unchanged in male neonates (diet: (*F*_(1,12)_ = 0.227, p = 0.643); treatment: (*F*_(3,12)_ = 0.658, p = 0.594)) (**Figure 6b**). Hsp70 protein abundance also remained unchanged in female neonates (diet: (*F*_(1,12)_ = 0.725, p = 0.411); treatment: (*F*_(3,12)_ = 1.181, p = 0.358)) (**Figure 6b**).

Hsp90 protein abundance remained unchanged in male neonates (diet: (*F*_(1,12)_ = 0.064, p = 0.805); treatment: (*F*_(3,12)_ = 0.043, p = 0.987) (**Figure 6c**). In female neonates, Hsp90 protein abundance was responsive to changes in diet (*F*_(1,12)_ = 10.665, p = 0.007). A treatment effect was observed (*F*_(3,12)_ = 3.561, p = 0.047), although no individual effects were observed between treatment groups (**Figure 6c**).

Hsp40 protein abundance in male neonates was responsive to diet (*F*_(1,12)_ = 58.828, p < 0.001), and MEV treatment (*F*_(3,12)_ = 20.299, p < 0.001), where Hsp40 protein abundance was higher in mCHD than mHFD (Tukey HSD, p < 0.001) and mHFD-MEV (Tukey HSD, p = 0.004) (**Figure 6d**). Similarly, Hsp40 protein abundance was higher in mCHD-MEV than mHFD (Tukey HSD, p < 0.001) and mHFD-MEV (Tukey HSD, p = 0.001). Hsp40 protein abundance in female neonates was also responsive to diet (*F*_(1,11)_ = 23.438, p < 0.001) and MEV treatment (*F*_(3,11)_ = 9.733, p = 0.002), where Hsp40 protein abundance was higher in mHFD-MEV than mCHD (Tukey HSD, p = 0.004) and mCHD-MEV (Tukey HSD, p = 0.004) (**Figure 6d**). Protein data of male and female neonates that received PBS gavage (the sham controls) is included in **Table S6**.

### 3.6 Transcript abundance of HSR targets in the hypothalamus

In the hypothalamus, HSF1 transcript abundance remained unchanged in male neonates (diet: (*F*_(1,12)_ = 3.954, p = 0.070); treatment: (*F*_(3,12)_ = 1.908, p = 0.182)) (**Figure 7a**). HSF1 transcript in female neonates was responsive to diet (*F*_(1,12)_ = 8.654, p = 0.012). The treatment effect approaches significance (*F*_(3,12)_ = 3.465, p = 0.051) (**Figure 7a**).

**Figure 7.**
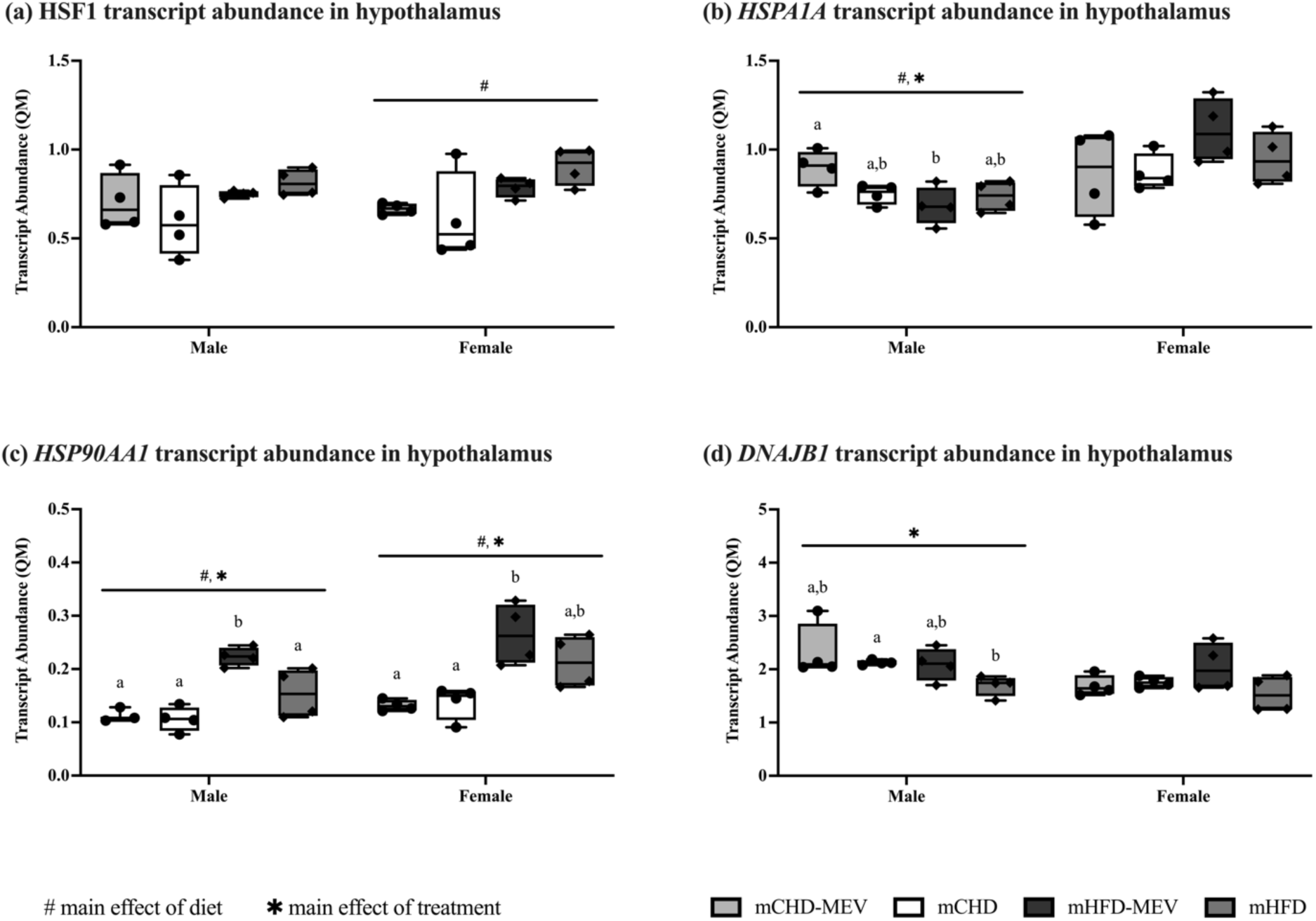
Transcript abundance of the heat shock response genes in the hypothalamus of male and female neonates at postnatal (PND) 11 as determined by RT-qPCR. (a) HSF1 hypothalamus transcript abundance. (b) *HSPA1A* hypothalamus transcript abundance. (c) *HSP90AA1* hypothalamus transcript abundance. (d) *DNAJB1* hypothalamus transcript abundance. Quantity means are normalized to one reference gene with stable expression: *GUSB*. ^#^Main effect of diet (p<0.05). *Main effect of MEV treatment (p<0.05). Pairwise comparisons between treatment groups are indicated with lowercase letters, where significant differences (p<0.05) are denoted by different letters. mCHD: Neonates born to mCHD dams that did not receive MEV supplementation. mCHD-MEV: Neonates born to mCHD dams that received MEV supplementation. mHFD: Neonates born to mHFD dams that did not receive MEV supplementation. mHFD-MEV: Neonates born to mHFD dams that received MEV supplementation. n = 3-4 biological replicates/diet/treatment/sex. Data presented are means ± standard error.

*HSPA1A* transcript abundance in male neonates was responsive to diet (*F*_(1,12)_ = 6.179, p = 0.029) and MEV treatment (*F*_(3,12)_ = 4.079, p = 0.033) where *HSPA1A* transcript abundance was higher in mCHD-MEV than mHFD-MEV (Tukey HSD, p = 0.026) (**Figure 7b**). *HSPA1A* transcript in females remained unchanged (diet: (*F*_(1,12)_ = 3.343, p = 0.092); treatment (*F*_(3,12)_ = 1.642, p = 0.232)) (**Figure 7b**).

*HSP90AA1* transcript abundance in male neonates was responsive to diet (*F*_(1,11)_ = 27.483, p < 0.001) and MEV treatment (*F*_(3,11)_ = 13.174, p < 0.001), where *HSP90AA1* transcript was higher in mHFD-MEV than mCHD (Tukey HSD, p < 0.001), mCHD-MEV (Tukey HSD, p = 0.002), and mHFD (Tukey HSD, p = 0.028) (**Figure 7c**). *HSP90AA1* transcript abundance in female neonates was also responsive to diet (*F*_(1,12)_ = 25.717, p < 0.001), and MEV treatment (*F*_(3,12)_ = 9.613, p = 0.002), where *HSP90AA1* transcript was higher in mHFD-MEV than mCHD (Tukey HSD, p = 0.004) and mCHD-MEV (Tukey HSD, p = 0.003) (**Figure 7c**).

*DNAJB1* transcript abundance in male neonates did not change with diet (*F*_(1,11)_ = 4.272, p = 0.063), although a treatment effect was observed (*F*_(3,11)_ = 4.364, p = 0.030) where *DNAJB1* transcript was higher in mCHD than mHFD (Tukey HSD, p = 0.039) (**Figure 7d**). *DNAJB1* transcript remained unchanged in female neonates (diet: (*F*_(1,12)_ = 0.210, p = 0.655); treatment (*F*_(3,12)_ = 1.988, p = 0.170)) (**Figure 7d**). Transcript data of male and female neonates that received PBS gavage (the sham controls) is included in **Table S6**.

### 3.7 Protein abundance of HSR targets in the hypothalamus

In the hypothalamus, HSF1 protein abundance in male neonates remained unchanged (diet: (*F*_(1,10)_ = 1.627, p = 0.231); treatment (*F*_(3,10)_ = 0.985, p = 0.438)) (**Figure 8a**). HSF1 protein abundance in female neonates was responsive to diet (*F*_(1,10)_ = 10.413, p = 0.009) and MEV treatment (*F*_(3,10)_ = 4.180, p = 0.037), where HSF1 protein was higher in mHFD than mCHD (Tukey HSD, p = 0.036) (**Figure 8a**).

**Figure 8.**
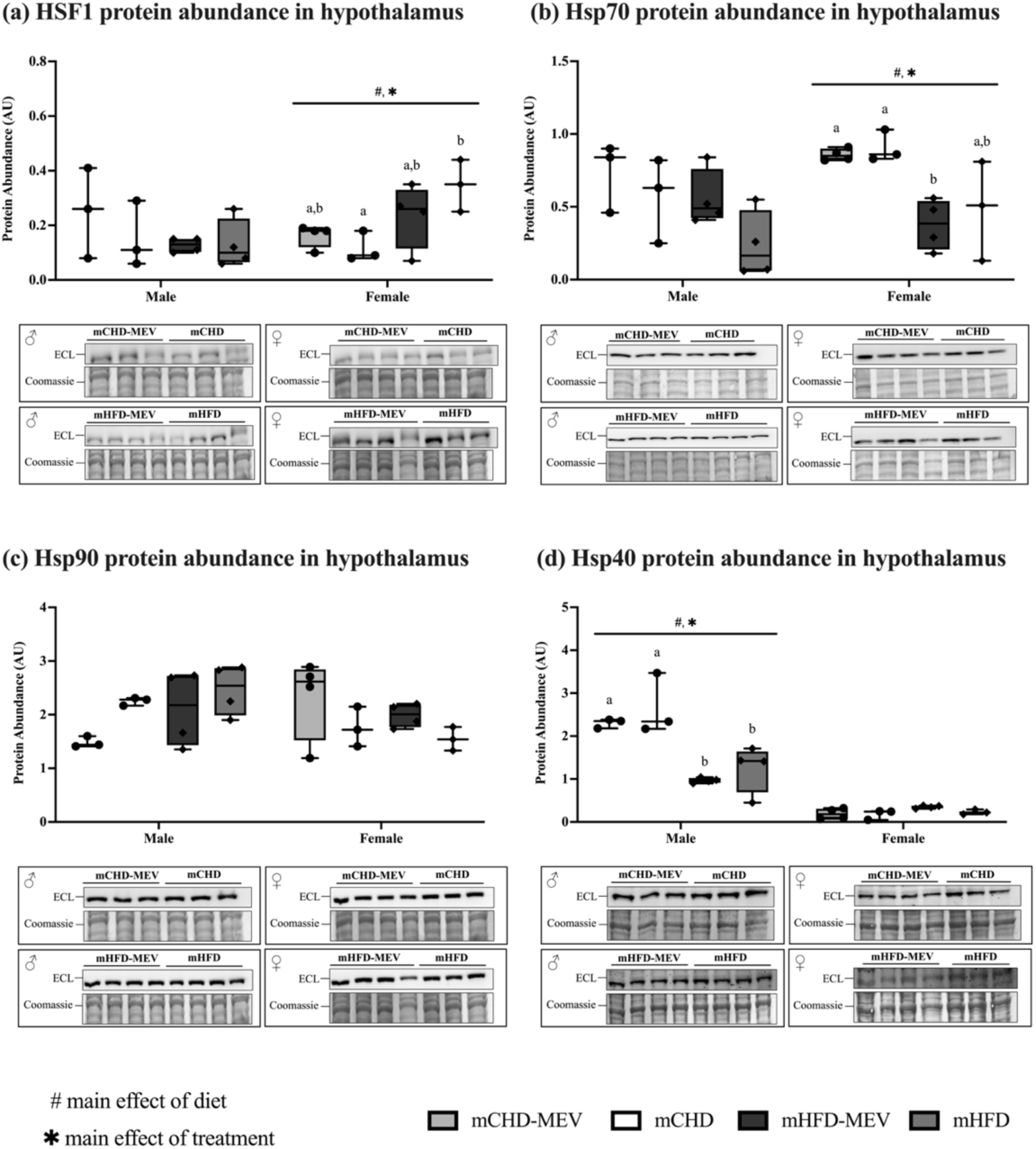
Protein abundance of the heat shock response targets in the hypothalamus of male and female neonates at postnatal day (PND) 11 as determined by western immunoblotting. **(a)** HSF1 hypothalamus protein abundance. **(b)** Hsp70 hypothalamus protein abundance. **(c)** Hsp90 hypothalamus protein abundance. **(d)** Hsp40 hypothalamus protein abundance. Protein targets are normalized to the abundance of total soluble proteins in the samples using Coomassie staining. ECL and Coomassie-stained images are displayed. ^#^Main effect of diet (p<0.05). *Main effect of treatment (p<0.05). Pairwise comparisons between treatment groups are indicated with lowercase letters, where significant differences (p<0.05) are denoted by different letters. mCHD: Neonates born to mCHD dams that did not receive MEV supplementation. mCHD-MEV: Neonates born to mCHD dams that received MEV supplementation. mHFD: Neonates born to mHFD dams that did not receive MEV supplementation. mHFD-MEV: Neonates born to mHFD dams that received MEV supplementation. n = 3-4 biological replicates/diet/treatment/sex. Data presented are means ± standard error.

Hsp70 protein abundance in male neonates remained unchanged (diet: (*F*_(1,10)_ = 3.987, p = 0.074); treatment (*F*_(3,10)_ = 2.835, p = 0.092)) (**Figure 8b**). Hsp70 protein abundance in female neonates was responsive to diet (*F*_(1,10)_ = 20.403, p = 0.001) and MEV treatment (*F*_(3,10)_ = 7.324, p = 0.007) where Hsp70 protein was lower in mHFD-MEV than mCHD (Tukey HSD, p = 0.016) and mCHD-MEV (Tukey HSD, p = 0.019) (**Figure 8b**).

Hsp90 protein abundance in male neonates remained unchanged (diet: (*F*_(1,10)_ = 2.687, p = 0.132); treatment (*F*_(3,10)_ = 2.647, p = 0.106)) (**Figure 8c**). Hsp90 protein abundance in female neonates also remained unchanged (diet: (*F*_(1,10)_ = 1.162, p = 0.306); treatment (*F*_(3,10)_ = 1.686, p = 0.232)) (**Figure 8c**).

Hsp40 protein abundance in male neonates was responsive to diet (*F*_(1,10)_ = 33.566, p < 0.001), and MEV treatment (*F*_(3,10)_ = 11.790, p = 0.001), where Hsp40 protein was higher in mCHD compared to mHFD (Tukey HSD, p = 0.008) and mHFD-MEV (Tukey HSD, p = 0.002). Hsp40 protein was also higher in mCHD-MEV compared to mHFD (Tukey HSD, p = 0.043) and mHFD-MEV (Tukey HSD, p = 0.011) (**Figure 8d**). Hsp40 protein abundance in female neonates remained unchanged (diet: (*F*_(1,10)_ = 4.794, p = 0.053); treatment (*F*_(3,10)_ = 2.847, p = 0.091) (**Figure 8d**). Protein data of male and female neonates that received PBS gavage (the sham controls) is included in **Table S6**.

### 3.8 Transcript abundance of HSR targets in the prefrontal cortex

In the prefrontal cortex, HSF1 transcript abundance in male neonates remained unchanged (diet: (*F*_(1,12)_ = 2.867, p = 0.116); treatment (*F*_(3,12)_ = 1.486, p = 0.268)) (**Figure 9a**). HSF1 transcript abundance in female neonates also remained unchanged (diet: (*F*_(1,11)_ = 0.902, p = 0.363); treatment (*F*_(3,11)_ = 0.920, p = 0.463)) (**Figure 9a**).

**Figure 9.**
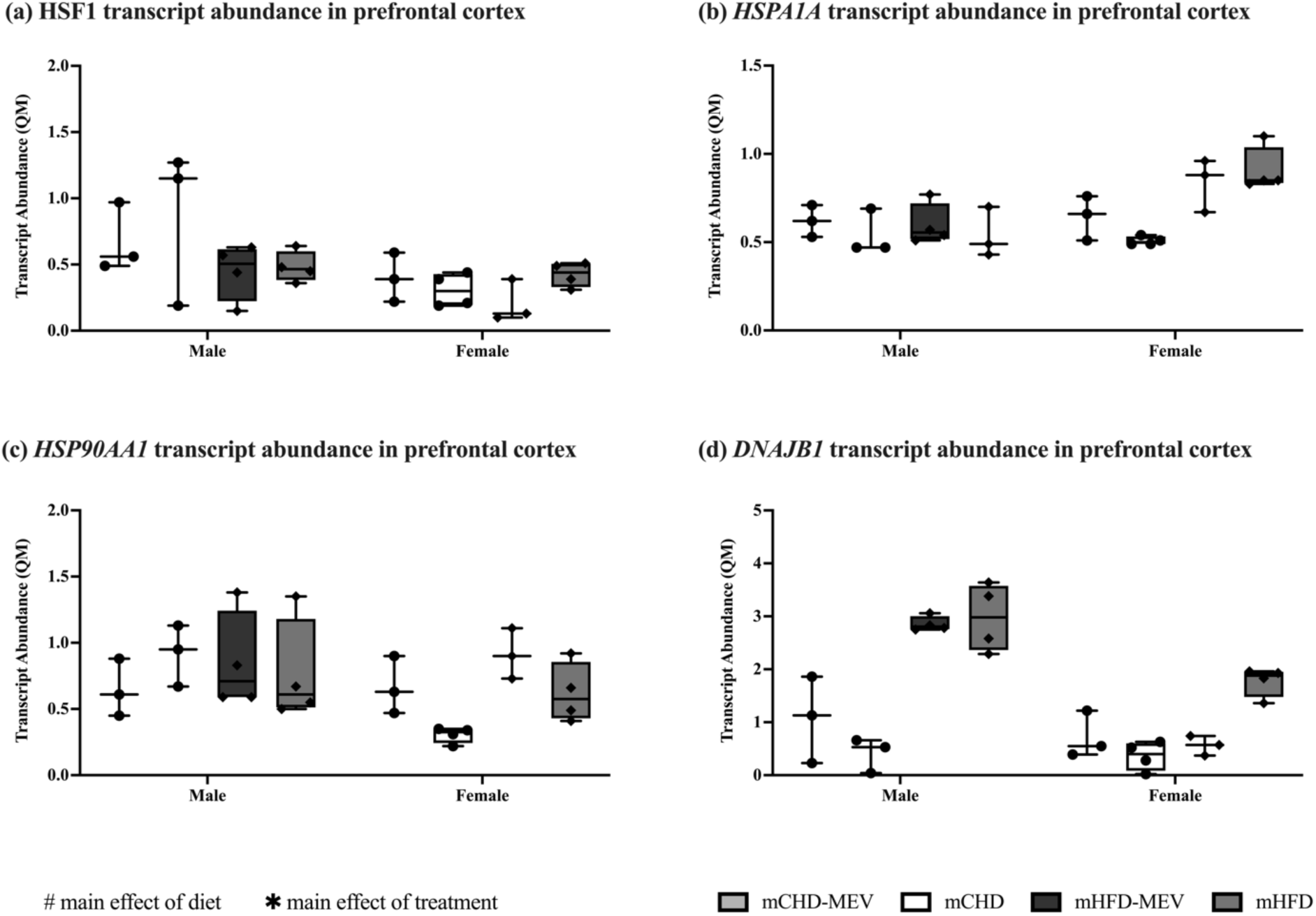
Transcript abundance of the heat shock response genes in the prefrontal cortex of male and female neonates at postnatal (PND) 11 as determined by RT-qPCR. **(a)** HSF1 prefrontal cortex transcript abundance. **(b)** *HSPA1A* prefrontal cortex transcript abundance. **(c)** *HSP90AA1* prefrontal cortex transcript abundance. **(d)** *DNAJB1* prefrontal cortex transcript abundance. Quantity means are normalized to the geometric mean of two reference genes with stable expression: *18S* rRNA and *YWAZ*. ^#^Main effect of diet (p<0.05). *Main effect of MEV treatment (p<0.05). Pairwise comparisons between treatment groups are indicated with lowercase letters, where significant differences (p<0.05) are denoted by different letters. mCHD: Neonates born to mCHD dams that did not receive MEV supplementation. mCHD-MEV: Neonates born to mCHD dams that received MEV supplementation. mHFD: Neonates born to mHFD dams that did not receive MEV supplementation. mHFD-MEV: Neonates born to mHFD dams that received MEV supplementation. n = 3-4 biological replicates/diet/treatment/sex. Data presented are means ± standard error.

*HSPA1A* transcript abundance in male neonates remained unchanged (diet: (*F*_(1,12)_ = 0.565, p = 0.467); treatment (*F*_(3,12)_ = 1.918, p = 0.181)) (**Figure 9b**). *HSPA1A* transcript abundance also remained unchanged in female neonates (diet: (*F*_(1,11)_ = 1.416, p = 0.259); treatment (*F*_(3,11)_ = 1.016, p = 0.423)) (**Figure 9b**).

*HSP90AA1* transcript abundance in male neonates remained unchanged (diet: (*F*_(1,12)_ = 0.717, p = 0.414); treatment (*F*_(3,12)_ = 0.328, p = 0.805)) (**Figure 9c**). *HSP90AA1* transcript abundance also remained unchanged in female neonates (diet: (*F*_(1,12)_ = 0.010, p = 0.923); treatment (*F*_(3,12)_ = 0.044, p = 0.987)) (**Figure 9c**).

*DNAJB1* transcript abundance in male neonates remained unchanged (diet: (*F*_(1,12)_ = 2.620, p = 0.131); treatment (*F*_(3,12)_ = 2.537, p = 0.106)) (**Figure 9d**). *DNAJB1* transcript abundance also remained unchanged in female neonates (diet: (*F*_(1,12)_ = 4.292, p = 0.060); treatment (*F*_(3,12)_ = 3.002, p = 0.073)) (**Figure 9d**). Transcript data of male and female neonates that received PBS gavage (the sham controls) is included in **Table S6**.

### 3.9 Protein abundance of HSR targets in the prefrontal cortex

In the prefrontal cortex, HSF1 protein abundance remained unchanged in male neonates (diet: (*F*_(1,10)_ = 0.532, p = 0.483); treatment (*F*_(3,10)_ = 2.634, p = 0.107)) (**Figure 10a**). HSF1 protein abundance in female neonates did not change with diet (*F*_(1,10)_ = 2.414, p = 0.151), but there was an effect of treatment (*F*_(3,10)_ = 6.884, p = 0.009), where HSF1 protein was higher in mHFD-MEV compared to mCHD-MEV (Tukey HSD, p = 0.008) and mHFD (Tukey HSD, p = 0.044) (**Figure 10a**).

**Figure 10.**
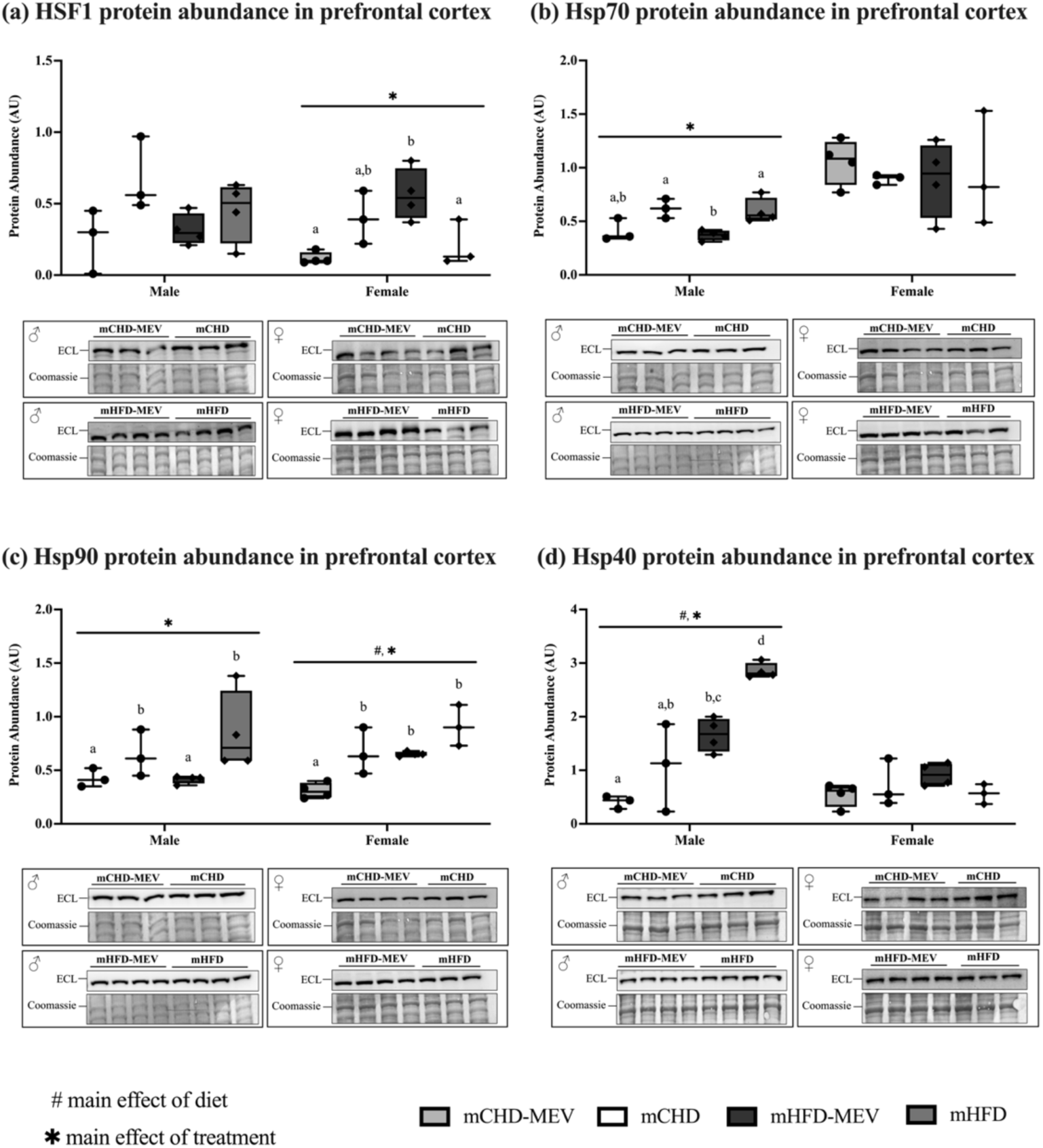
Protein abundance of the heat shock response targets in the prefrontal cortex of male and female neonates at postnatal day (PND) 11 as determined by western immunoblotting. **(a)** HSF1 prefrontal cortex protein abundance. **(b)** Hsp70 prefrontal cortex protein abundance. **(c)** Hsp90 prefrontal cortex protein abundance. **(d)** Hsp40 prefrontal cortex protein abundance. Protein targets are normalized to the abundance of total soluble proteins in the samples using Coomassie staining. ECL and Coomassie-stained images are displayed. ^#^Main effect of diet (p<0.05). *Main effect of treatment (p<0.05). Pairwise comparisons between treatment groups are indicated with lowercase letters, where significant differences (p<0.05) are denoted by different letters. mCHD: Neonates born to mCHD dams that did not receive MEV supplementation. mCHD-MEV: Neonates born to mCHD dams that received MEV supplementation. mHFD: Neonates born to mHFD dams that did not receive MEV supplementation. mHFD-MEV: Neonates born to mHFD dams that received MEV supplementation. n = 3-4 biological replicates/diet/treatment/sex. Data presented are means ± standard error.

Hsp70 protein abundance in male neonates did not change with diet (*F*_(1,10)_ = 0.383, p = 0.550) but there was an effect of treatment (*F*_(3,10)_ = 6.703, p = 0.009), where Hsp70 protein was lower in mHFD-MEV than mCHD (Tukey HSD, p = 0.024) and mHFD (Tukey HSD, p = 0.026) (**Figure 10b**). Hsp70 protein abundance remained unchanged in female neonates (diet: (*F*_(1,10)_ = 0.096, p = 0.763); treatment (*F*_(3,10)_ = 0.208, p = 0.889) (**Figure 10b**).

Hsp90 protein abundance in male neonates did not change with diet (*U* = 20.000, p = 0.945), but there was a treatment effect (*H*(3) = 8.170, p = 0.043), where MEV supplementation reduced the protein levels of Hsp90 compared to the respective controls. Specifically, mCHD-MEV was lower than mCHD (Dunn’s post hoc, p = 0.046) and mHFD (Dunn’s post hoc, p = 0.046). Similarly, mHFD-MEV was lower than mCHD (Dunn’s post hoc, p = 0.041) and mHFD (Dunn’s post hoc, p = 0.041) (**Figure 10c**). Hsp90 protein abundance in female neonates was responsive to diet (*F*_(1,10)_ = 16.320, p = 0.002), and MEV treatment (*F*_(3,10)_ = 11.946, p = 0.001), where Hsp90 protein was lower in mCHD-MEV than mCHD (Tukey HSD, p = 0.027), mHFD (Tukey HSD, p < 0.001), and mHFD-MEV (Tukey HSD, p = 0.022) (**Figure 10c**).

Hsp40 protein abundance in male neonates was responsive to diet (*F*_(1,10)_ = 45.741, p < 0.001), and MEV treatment (*F*_(3,10)_ = 22.029, p < 0.001) where Hsp40 protein was lower in mCHD-MEV than mHFD-MEV (Tukey HSD, p = 0.012) (**Figure 10d**). Additionally, Hsp40 protein was higher in mHFD than mCHD (Tukey HSD, p = 0.001), mCHD-MEV (Tukey HSD, p < 0.001) and mHFD-MEV (Tukey HSD, p = 0.010). Hsp40 protein abundance in female neonates remained unchanged (diet: (*F*_(1,10)_ = 0.559, p = 0.472); treatment (*F*_(3,10)_ = 1.585, p = 0.254)) (**Figure 10d**). Protein data of male and female neonates that received PBS gavage (the sham controls) is included in **Table S6**.

## 4.0 Discussion

We investigated the interactions between MEVs and the HSR in the liver, hypothalamus, and prefrontal cortex in male and female neonatal rats exposed to mHFD and mCHD at PND11. PND11 is a critical window of early development in Long Evans rats and represents the peak lactational period and it is within the SHRP that spans the first 14 days of postnatal life (Burnol et al., 1983; Jakubowski and Terkel, 1986; Sharma et al., 2022). Although acute stress that leads to heightened glucocorticoid release is reduced during SHRP (Sapolsky and Meaney, 1986), chronic stress such as mHFD, leads to significant developmental and inflammatory outcomes in offspring (Abuaish et al., 2018; Sasaki et al., 2014, 2013; Wijenayake et al., 2020). The HSP chaperones involved in the HSR participate in protein folding and refolding mechanisms, disaggregation, and degradation (Hu et al., 2022). Hsp70 and Hsp90 form the Hsp70-Hsp90 complex, a critical cellular machinery for protein refolding and disaggregation, which is assisted by co-chaperone Hsp40 to identify and deliver misfolded proteins to the complex (Doyle and Wang, 2019; Hernández et al., 2002; Qiu et al., 2006). The gene expression of these chaperones is regulated by the primary transcription factor, HSF1 (Jaeger et al., 2014). Our overall findings indicated target, tissue, and sex-specific interactions between MEVs and the HSR, where MEV treatment to offspring with perinatal mHFD upregulated HSF1 expression and downregulated the expression of its inhibitors in the prefrontal cortex. Further, females were more responsive to MEV treatment than male littermates.

We observed minimal differences in bodyweight between mCHD and mHFD dams during pre-gestation, gestation, and lactation (**Figure 2a**). Previous studies have reported that dams on this particular high fat diet tend to exhibit higher bodyweights than mCHD dams (Abuaish et al., 2018; Sasaki et al., 2013). However, these studies used a rodent chow as the control diet and did not use the same mCHD as our study, which is matched in sucrose content to the mHFD. Body weight gain is directly proportional to the dietary composition and palatability in rats with diet-induced obesity (Levin and Dunn-Meynell, 2002). Nevertheless, as shown by Abuaish et al. (2018) and Sasaki et al. (2013), the mHFD dams consumed more kcal than mCHD dams during pre-gestation (**Figure 2e**).

Our offspring data also corroborates previous studies that have used the same mHFD to induce perinatal diet stress in Long Evans neonates (Abuaish et al., 2018; Sasaki et al., 2013), where mHFD offspring were significantly heavier than mCHD offspring postnatally (**Figure 3a**).The Lee index, a measure of obesity in rodents that accounts for naso-anal length and bodyweight (Simson and Gold, 1982), was also greater in mHFD offspring compared to mCHD offspring (**Figure 3b**). Correspondingly, at PND11, mHFD offspring had significantly higher amounts of retroperitoneal fat than mCHD offspring, as well as significantly higher amounts of stomach milk curd (**Figure 4a/c)**. In combination, our data shows that mHFD exposure during perinatal life leads to increased bodyweight, Lee Index, and adiposity in both males and female neonates and these changes occur during the SHRP. We did not find sex or treatment effects in bodyweight and Lee Index.

The liver is a multicellular organ that plays significant roles in detoxification, hematopoiesis, protein synthesis, metabolism, bile production, and regulation of the innate immune system (Ishibashi et al., 2009; Liu and Yin, 2022). Multiple studies have reported that small EVs bioaccumulate in the liver following administration, before being repackaged and re-distributed to other organs (Kang et al., 2021; Manca et al., 2018; Yáñez-Mó et al., 2015; Zhang et al., 2020, 2016). Furthermore, the liver is also impacted by pro-inflammation via the liver-gut-brain axis (Butterworth, 2013), in particular with perinatal mHFD exposure (van der Heijden et al., 2015; Huang et al., 2017). As such, we investigated if and how MEVs may regulate the HSR in neonatal liver in response to chronic diet stress. We found a strong sex effect in HSR at the transcript and protein levels, where HSR in females was more responsive compared to male littermates. These findings are consistent with previous studies that have found females to be more sensitive to perinatal nutritional stress than males during the SHRP (Abuaish et al., 2018; Sasaki et al., 2013; Wijenayake et al., 2020). At the transcript level, females with perinatal mCHD responded to MEV treatment more robustly than males, with increased HSF1, *HSPA1A*, and *DNAJB1* (**Figure 5a/b/d).** No changes were seen at the protein level across sex, diet, and treatment (**Figure 6**), with exception to Hsp40, where mHFD males had lower Hsp40, and mHFD females had higher Hsp40 (**Figure 6d**). Further studies are required to determine if this inverse regulation in Hsp40 between sexes influences overall cytoprotection and capability to withstand and respond to proteotoxic stress in early life. In addition, transcript level changes in HSR in mCHD offspring did not correlate with protein abundance. It is possible that these responses are not being translated to the protein level because the baseline protein levels of the candidate HSPs are sufficient to combat proteotoxic stress and additional increases in cytoprotection are not required. Distribution patterns and expression of Hsp70 and Hsp40 protein and RNA (Karlsson et al., 2021; Sjöstedt et al., 2020; Thul et al., 2017; Uhlén et al., 2015), summarized in the Human Protein Atlas (proteinatlas.org), are typically shown to be higher in the liver compared to the CNS. Specifically, Hsp70 has higher protein levels in cholangiocytes and hepatocytes compared to lower protein levels detected in endothelial, glial and neuronal cells in the cerebral cortex. *HSPA1A* abundance in the liver is 1089.4 normalized transcripts per million (nTPM) compared to 442.9 nTPM in the cerebral cortex. While Hsp40 has similar protein levels in cholangiocytes and hepatocytes compared to cerebral cortex endothelial and glial cells, *DNAJB1* abundance is higher in the liver (224.7 nTPM) than the cerebral cortex (157.8 nTPM). Protein levels of HSF1 are conversely found to be higher in the brain than the liver, but RNA levels remain similar between the liver (44.9 nTPM) and cerebral cortex (46.5 nTPM). Therefore, it is possible that the endogenous HSR is sufficient in providing necessary levels of cytoprotection, so that MEV treatment does not produce additional effects.

An alternative explanation for the lack of changes with MEV treatment in the liver could be due to the timing of EV uptake by the liver. Several studies have described the liver as an active site of EV uptake (Szabo and Momen-Heravi, 2017). A study by Bala et al. (2015) which delivered miR-155-enriched B-cell derived EVs to miR-155 deficient C57/Bl6J mice via intraperitoneal or intravenous injections, found that the enriched EVs rapidly biodistributed with the highest amount detected in the liver, as rapidly as within 10 minutes. Similarly, a study by Manca et al. (2018) found that concentrations of bovine MEVs administered to Balb/c mice were highest in the liver and spleen, while the biodistribution of MEV miRNA cargo was dissimilar, with majority localizing to the brain. Furthermore, Manca et al. (2018) corroborated a rapid uptake of EVs in the liver, as MEV concentrations peaked 3h after intravenous injection and 12h after administration via oral gavage. Together these studies suggest that EVs are rapidly bioaccumulated in the liver, where the cargo may be repackaged from exogenous EVs to endogenous EVs and redistributed. It is possible that with our results, the reason we do not observe significant responses to MEV treatment is because EV uptake typically occurs within the first 12h, while this study was conducted using tissues collected at minimum 16h after the last MEV treatment.

The hypothalamus is responsible for thermoregulation, circadian rhythm, feeding behaviour, energy metabolism, and social responses and is a central component of the limbic system (Coll and Yeo, 2013; Williams et al., 2001). Furthermore, the PVN of the hypothalamus directly regulates HPA axis activity (Niu et al., 2019). At the transcript level in the hypothalamus, no changes were seen with MEV treatment across sex, diet, or treatment (**Figure 7a-d**), with exception to *HSP90AA1* in males, which decreased with MEV treatment (**Figure 7c**). At the protein level, no changes were seen with MEV across sex, diet, or treatment (**Figure 8a-d**). The only significant effects seen in the hypothalamus were diet-induced, where, at the transcript level, mHFD increased HSF1 and *HSP90AA1*, while decreasing *HSPA1A* (**Figure 7a-c**). Diet effects at the protein level were also observed where mHFD increased HSF1 and decreased Hsp70 and Hsp40 (**Figure 8a/b/d**). Given that the hypothalamus plays a central role in regulating feeding behaviour, HPA axis regulation and is sensitive to diet-induced pro-inflammation, it is unsurprising that we primarily see diet effects. A study by De Souza et al. (2005) analyzed the hypothalamus of male Wistar rats following 16-week consumption of a standard diet (10% kcal lipids) or hyperlipidic diet (45% kcal lipids). They found that a hyperlipidic diet induces the expression of several pro-inflammatory proteins including TNFα, IL-1β, and IL-6. Furthermore, they also reported that a hyperlipidic diet results in increased c-Jun N-terminal kinase and NFκB activation, corroborating the sensitivity of the hypothalamus to diet stress. Likely diet effects resulted in heightened baseline levels in mHFD exposed offspring, and MEVs were unable to exert strong cytoprotective effects. Furthermore, the hypothalamus is a heterogeneous brain region with many, distinct nuclei and subnuclei which are spatially separated into the preoptic, tuberal and posterior regions (Huang et al., 2024; Saper and Lowell, 2014). Given its heterogeneity, it is likely that MEV effects are difficult to delineate in the whole hypothalamus, as opposed to distinct nuclei within the hypothalamus, such as the PVN. A study by Huang et al. (2024) analyzed single nucleus transcriptomics in the hypothalamus of PND15 C57BL/6 mice exposed to mCHD (10% kcal fat) or mHFD (45% kcal fat). They found that within the 30 cellular subpopulations analyzed within the hypothalamus, that neuronal subpopulations respond to mHFD differentially, especially for markers involved in feeding behaviour, glucose homeostasis, insulin signaling, and circadian rhythm. This further supports the difficulty in assessing the effects of MEV treatment in response to mHFD in the hypothalamus, rather than assessing individual hypothalamic nuclei.

The prefrontal cortex is involved in higher-order executive and cognitive function, forming extensive connections with other brain regions, including the hypothalamus and amygdala (Arnsten, 2009; Friedman and Robbins, 2022). It is involved in glucocorticoid receptor-mediated feedback inhibition of the HPA axis (Diorio et al., 1993; Laryea et al., 2015; McKlveen et al., 2013; Sullivan and Gratton, 2002). At the transcript level in the prefrontal cortex, MEV treatment did not lead to changes in HSR targets, irrespective of diet and sex (**Figure 9a-d**). At the protein level in the prefrontal cortex, protein abundance of HSR targets was not responsive to MEVs in mCHD neonates (**Figure 10a-d**), with exception to Hsp90 which decreased with MEV treatment in male and female neonates (**Figure 10c**). However, significant changes were in mHFD offspring at the protein level, where in mHFD males, protein abundance of Hsp70, Hsp90 and Hsp40 decreased with MEV treatment (**Figure 10b-d**). In mHFD female neonates, HSF1 increased with MEV treatment (**Figure 10a**). Hsp90 also decreased with MEV treatment, although this change was approaching statistical significance (**Figure 10c**). Previous studies corroborate our findings and report that increased HSF1 activity in the brain, including the prefrontal cortex and hippocampus, is neuroprotective against chronic stress (Fernandes et al., 2022; He et al., 2023; Wang et al., 2017). Furthermore, Hsp70 and Hsp90, members of the Hsp70-Hsp90 protein refolding complex, are implicated in the negative regulation of HSF1 (Prince et al., 2020). Hsp70 limits the hyperactivation and prolonged function of HSF1 by interfering with its binding and stabilization to keep it in a structurally inactive form (Prince et al., 2020). Concomitantly, Hsp90 disassembles structurally active HSF1, attenuating HSF1-mediated gene expression of HSP chaperones (Prince et al., 2020). Hsp40 acts as a co-chaperone that enhances the ATPase activity of Hsp70 and thus is typically found to follow the same regulatory patterns as Hsp70 (Eftekharzadeh et al., 2019; Michels et al., 1997). Our results illustrate that MEV treatment in the prefrontal cortex resulted in upregulation of HSF1, and downregulation of the negative regulators in male and female offspring with perinatal mHFD exposure. Furthermore, our results suggests that MEV treatment is prolonging the HSR, by maintaining HSF1 transcription. We reported a similar regulation *in vitro* using an immortalized human microglia clone 3 (HMC3) cell line, where MEV treatment increases protein abundance of HSF1, while decreasing the abundance of the Hsp70 and Hsp90 negative regulators (Storm et al., 2025).

The objective of this study was to explore if MEV treatment may modulate cytoprotection in response to perinatal mHFD exposure by interacting with the HSR in the liver, hypothalamus, and prefrontal cortex of male and female neonatal rats during the stress hyporesponsive period. In the liver, MEV treatment led to an upregulation of HSF1, *HSPA1A* and *DNAJB1* in female mCHD offspring, but the increase did not translate to protein abundance. This lack of correlation between transcript and protein levels in the liver may be due to fact that the baseline HSR is generally higher in the liver of neonates than in the hypothalamus and the prefrontal cortex, and a further upregulation of HSPs is not necessary to main homeostasis and combat proteotoxic stress. In the hypothalamus, the only significant effects observed were in response to maternal diet. In the prefrontal cortex, our results illustrate that MEV treatment exerted protective effects by influencing HSF1 regulation, prolonging its activation, while inducing the downregulation of the negative regulators, Hsp70, Hsp90, and co-chaperone Hsp40. To our knowledge, this study is the first to explore the relationship between MEVs and the HSR in male and female neonates in the context of promoting cytoprotection in the brain and the periphery in response to mHFD exposure. Now that we have established there are interactions between MEVs and the pro-survival HSR, further studies are warranted to determine associations between MEV cargo and these responses, and to elucidate the molecular mechanisms behind this control.

## 5.0 Conclusion

The therapeutic potential of MEVs increases as their role in improving the progression of various pro-inflammatory disease and afflictions is continually discovered. More recently, MEVs have been shown to cross the blood-brain barrier, delivering their cargo to regions within the brain. However, despite this knowledge, the cytoprotective potential of MEVs within the brain remains understudied. Our study is the first to investigate interactions between MEVs and pro-survival pathways in key tissues and regions involved in HPA axis-mediated stress responses, in the context of promoting cytoprotection. We found that the HSR was more robust in female neonates than males with MEV treatment. Through our perinatal mHFD model, we found that while MEV treatment only induces transcript-level changes in mCHD female offspring in the liver, and diet effects in the hypothalamus, robust effects are induced in the prefrontal cortex. Specifically, MEV supplementation increases the abundance of the main transcription factor HSF1, the main transcription factor, in the prefrontal cortex, while also downregulating its negative regulators, the Hsp70-Hsp90 complex, and Hsp70-binding partner Hsp40. We postulate that by downregulating Hsp70 and Hsp90, HSR activation is prolonged by MEV treatment, resulting in continued pro-survival benefits in response to mHFD stress. Future research will focus on investigating components of the MEV cargo inducing these effects as well as other downstream pro-survival targets regulated by HSF1 that may be activated.

## List of abbreviations

ACTH: Adrenocorticotrophic hormone
ATP: Adenosine triphosphate
AVP: Arginine vasopressin
BBB: Blood brain barrier
BCA: Bicinchoninic acid
cDNA: Complementary DNA
CNS: Central nervous system
CRH: Corticotrophin releasing hormone
DNA: Deoxyribonucleic acid
DOHaD: Developmental Origins of Health and Disease
ECL: Enhanecd chemiluminescence
EV: Extracellular vesicle
HFD: High fat diet
HPA: Hypothalamic-pituitary-adrenal
HSF: Heat shock transcription factor
HSP: Heat shock protein
HSR: Heat shock protein response
IBC: Interblot converter
mCHD: Perinatal control diet
MEV: Milk-derived extracellular vesicle
mHFD: Perinatal high fat diet
miRNA: MicroRNA
MISEV: Minimum information for studies of extracellular vesicles
MIQE: Minimum Information for Publication of Quantitative Real-time PCR
mRNA: Experiments Messenger RNA
MTT: 3-[4,5-dimethylthiazol-2-yl]-2,5 diphenyl tetrazolium bromide
NFκB: Nuclear factor kappa B
nTPM: Normalized transcripts per million
PBS: Phosphate-buffered saline
PND: Postnatal day
PVN: Paraventricular nucleus
RNA: Ribonucleic acid
RT-qPCR: Quantitative reverse transcription polymerase chain reaction
SDS: Sodium dodecylsulfate
SHRP: Stress hyporesponsive period

## Declarations

### Ethics approval

All animal studies complied with the Canadian Council on Animal Care Guidelines and Policies and were approved by the Local Animal Care Committee at the University of Winnipeg.

### Consent for publication

Not applicable.

### Availability of data and materials

All necessary datasets are included in the supplementary materials. Anything further can be provided upon request.

### Authors’ contributions

JAS was involved in data curation (related to animal care, tissue extractions, and gene and protein analysis of HSPs), formal analysis, investigation, validation, and was a major contributor in writing the manuscript (original draft, review, and editing). JL and MFO assisted in data curation related to animal experiments. SW was involved in conceptualization of the project, formal analysis, funding acquisition, supervision, and writing the manuscript (original draft, review, and editing). All authors read and approved the final manuscript.

## Acknowledgements

We would like to thank the support staff and vivarium staff at The University of Winnipeg, in particular Robyn Cole and Dan Wasyliw for assisting with the animal experiments. We would like to thank the research technicians at the Structural and Biophysical Core Facility at the Hospital for Sick Children (Toronto, ON, Canada) for conducting nanoparticle tracking analysis. We would like to thank the support staff at the Manitoba Institute for Materials at the University of Manitoba (Winnipeg, MB, Canada) for sharing their facilities and for use of the FEI Talos F200x S/TEM transmission electron microscope.

