## Supplementary Materials for "Human milk-derived extracellular vesicle treatment promotes the heat shock response in neonates with perinatal high fat diet exposure"

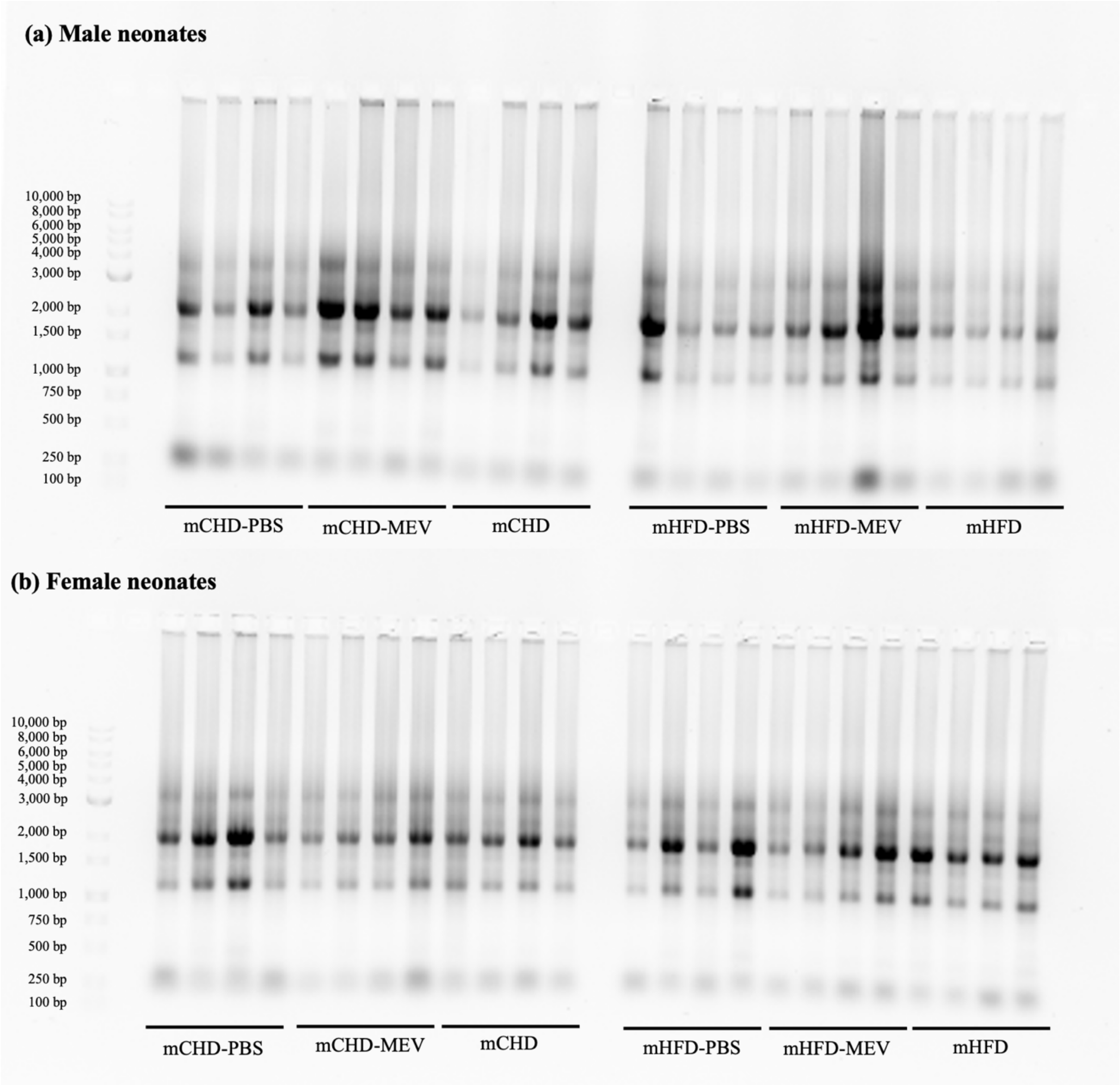


**Figure S1. RNA profiles for liver**. 1% TAE-agarose gel for verification of RNA stability and integrity. The ladder range is from 100 bp to 10,000 bp. RNA profiles for each group (mCHD-PBS, mCHD-MEV, mCHD, mHFD-PBS, mHFD-MEV, mHFD) show distinct 5S, 18S, and 28S subunit distribution.

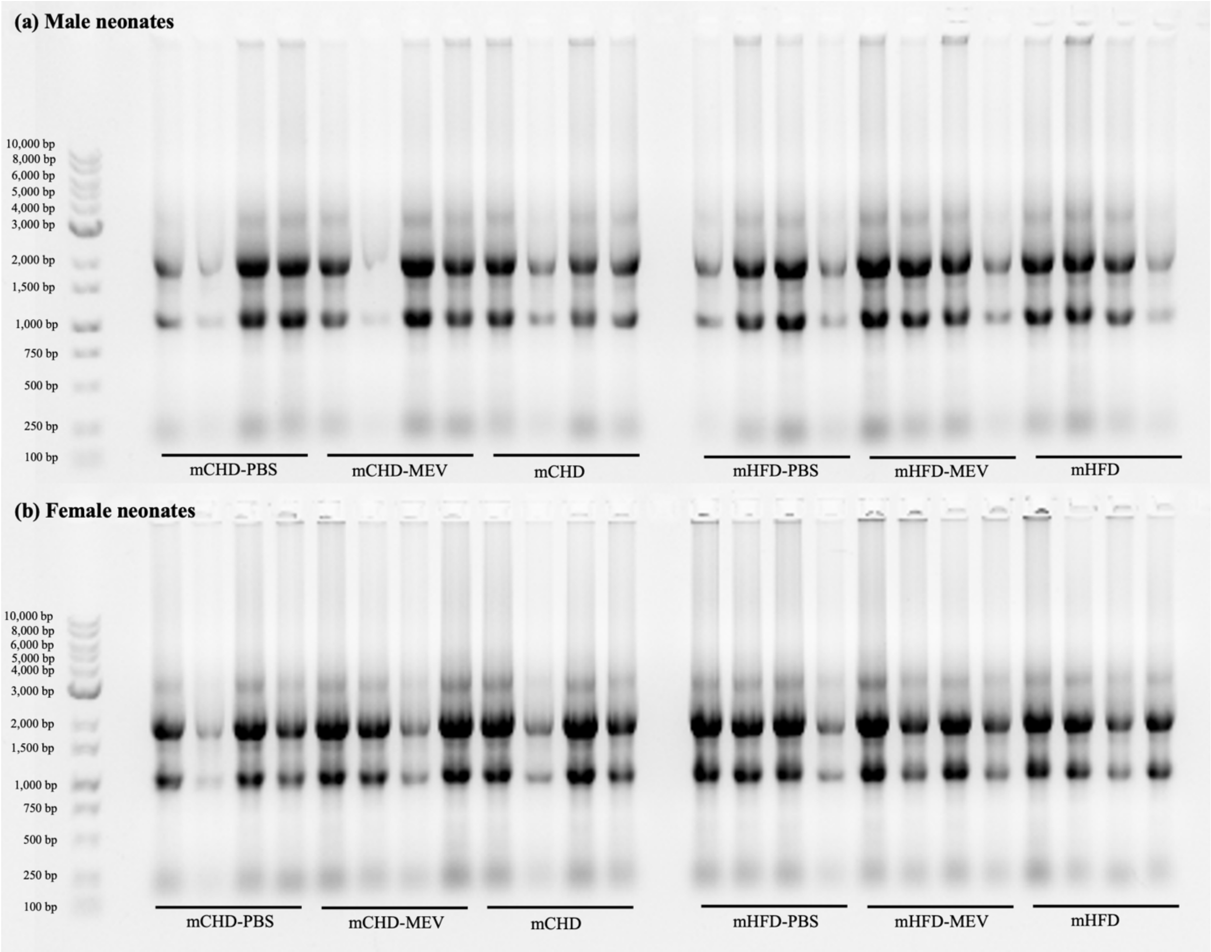


**Figure S2. RNA profiles for hypothalamus**. 1% TAE-agarose gel for verification of RNA stability and integrity. The ladder range is from 100 bp to 10,000 bp. RNA profiles for each group (mCHD-PBS, mCHD-MEV, mCHD, mHFD-PBS, mHFD-MEV, mHFD) show distinct 5S, 18S, and 28S subunit distribution.

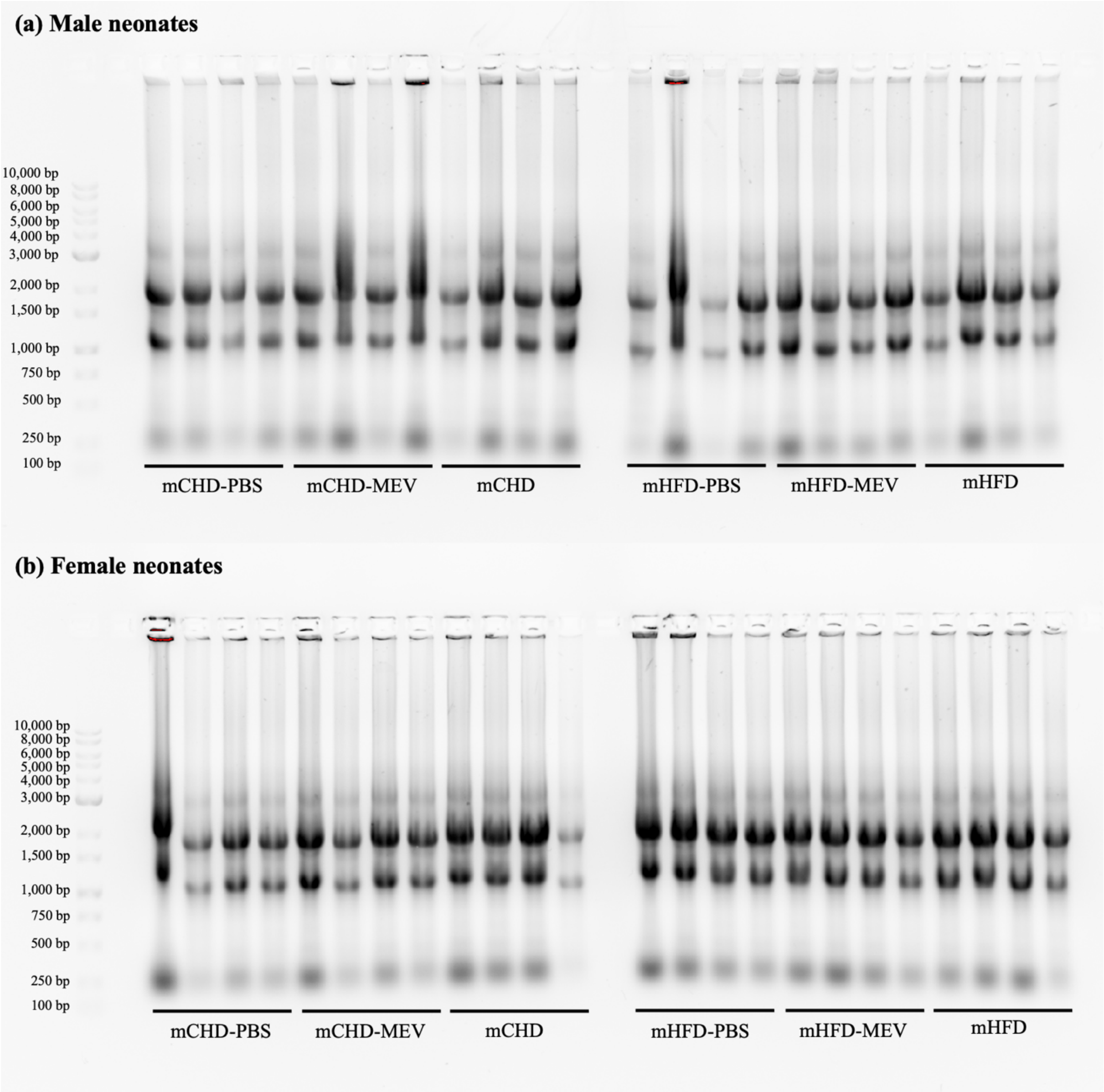


**Figure S3. RNA profiles for prefrontal cortex**. 1% TAE-agarose gel for verification of RNA stability and integrity. The ladder range is from 100 bp to 10,000 bp. RNA profiles for each group (mCHD-PBS, mCHD-MEV, mCHD, mHFD-PBS, mHFD-MEV, mHFD) show distinct 5S, 18S, and 28S subunit distribution.

**Table S1. Transcript data for candidate internal control genes that were tested in the liver, hypothalamus, and prefrontal cortex but were determined to be unsuitable for normalization.** The data are organized by tissue, target, sex, and treatment group. Transcript data are presented as quantity means, with the mean and SEM indicated for each treatment group. The main effects of treatment and diet are indicated (p<0.05).

| Tissue | Target | Sex | mCHD-PBS | | mCHD-MEV | | mCHD | | mHFD-PBS | | mHFD-MEV | | mHFD | | p-value | |
|---|---|---|---|---|---|---|---|---|---|---|---|---|---|---|---|---|
| | | | Mean | SEM | Mean | SEM | Mean | SEM | Mean | SEM | Mean | SEM | Mean | SEM | Treatment | Diet |
| Liver | *18S rRNA* | F | 9.98 | 1.65 | 8.12 | 0.52 | 11.19 | 1.99 | 6.44 | 0.13 | 7.38 | 0.43 | 6.81 | 0.47 | 0.839 | 0.004 |
| | *Actin-β* | M | 7.19 | 0.42 | 10.31 | 1.64 | 10.82 | 1.54 | 6.01 | 0.86 | 5.60 | 1.27 | 4.00 | 0.13 | 0.491 | <0.001 |
| Hypo-thalamus | *18S rRNA* | F | 9.59 | 0.64 | 9.38 | 0.98 | 8.37 | 0.60 | 18.00 | 1.99 | 12.66 | 0.70 | 9.79 | 0.76 | <0.001 | <0.001 |
| | *Actin-β* | F | 7.68 | 0.96 | 8.65 | 1.39 | 7.61 | 0.62 | 13.97 | 2.15 | 12.30 | 1.94 | 10.50 | 2.61 | 0.158 | 0.021 |
| | *B2M* | F | 6.19 | 0.09 | 6.32 | 0.65 | 5.00 | 0.48 | 10.60 | 1.13 | 9.30 | 1.16 | 8.20 | 1.58 | 0.203 | <0.001 |
| | *Eif1a1* | F | 6.66 | 0.71 | 6.64 | 0.74 | 5.36 | 0.64 | 11.41 | 1.57 | 11.53 | 1.41 | 8.62 | 1.92 | 0.196 | <0.001 |
| | *GAPDH* | F | 7.89 | 0.57 | 8.50 | 0.55 | 7.63 | 0.39 | 14.40 | 0.86 | 11.66 | 1.36 | 8.88 | 1.21 | 0.016 | <0.001 |
| | *GDI1* | F | 7.59 | 0.66 | 8.47 | 0.78 | 6.55 | 0.46 | 13.74 | 0.97 | 13.57 | 1.60 | 10.32 | 1.73 | 0.073 | <0.001 |
| | *Nupl2* | M | 38.93 | 5.26 | 56.42 | 2.88 | 56.57 | 4.24 | 54.10 | 16.92 | 66.58 | 8.63 | 79.28 | 4.97 | 0.060 | 0.034 |
| | *PGC1α* | F | 6.76 | 0.33 | 7.77 | 0.66 | 7.10 | 0.76 | 12.55 | 1.34 | 12.36 | 1.49 | 9.52 | 1.81 | 0.324 | <0.001 |
| | *PPIA* | F | 7.30 | 0.34 | 8.09 | 0.49 | 5.68 | 0.41 | 12.63 | 0.74 | 11.56 | 1.22 | 9.00 | 1.26 | 0.008 | <0.001 |
| | *Rpl5* | M | 4.86 | 0.66 | 7.19 | 1.14 | 8.15 | 0.79 | 12.67 | 2.02 | 11.59 | 0.76 | 10.13 | 0.69 | 0.856 | <0.001 |
| | *Rpl27* | F | 7.50 | 0.88 | 7.42 | 0.95 | 6.00 | 0.63 | 12.71 | 1.47 | 11.39 | 1.27 | 9.77 | 1.78 | 0.207 | <0.001 |
| | *Rpl32* | F | 7.70 | 0.23 | 7.26 | 0.75 | 5.25 | 0.64 | 12.39 | 1.46 | 9.88 | 0.54 | 7.59 | 1.38 | 0.004 | <0.001 |
| | *SDHA* | F | 8.59 | 0.67 | 9.32 | 0.75 | 7.66 | 0.56 | 15.90 | 1.77 | 14.81 | 2.57 | 10.43 | 2.00 | 0.104 | <0.001 |
| | *TBP* | F | 5.41 | 0.36 | 5.50 | 0.85 | 4.33 | 0.69 | 9.37 | 1.54 | 9.49 | 1.37 | 7.73 | 1.74 | 0.396 | 0.002 |
| | *YWAZ* | F | 7.44 | 0.73 | 9.36 | 0.75 | 7.82 | 0.55 | 15.20 | 1.75 | 13.10 | 1.80 | 10.21 | 1.42 | 0.151 | <0.001 |
| Prefrontal Cortex | *Actin-β* | F | 5.35 | 0.59 | 4.69 | 0.43 | 4.71 | 0.38 | 6.00 | 0.47 | 6.37 | 0.34 | 6.34 | 0.47 | 0.932 | 0.002 |
| | *GAPDH* | F | 5.60 | 0.15 | 5.27 | 0.19 | 5.32 | 0.21 | 5.97 | 0.43 | 5.79 | 0.40 | 6.56 | 0.34 | 0.419 | 0.010 |

**Table S2. Primer sequences and RT-qPCR parameters for HSR genes in the rat model**. Targets are identified by gene name, NCBI accession number, forward and reverse primer sequences, annealing temperature (°C), cDNA amount (ng), and their protein product. The sources of the primers (designed using NCBI or obtained from literature) are indicated. All primers target *Rattus norvegicus*.

| Gene | NCBI Accession # | Forward primer (5'–3') | Reverse primer (5'–3') | Annealing temperature (°C) | Liver cDNA loaded (ng) | Hypothalamus cDNA loaded (ng) | Prefrontal cortex cDNA loaded (ng) | Primer source |
|---|---|---|---|---|---|---|---|---|
| *18S rRNA* | NR_046237.3 | ATGGTAGTCGC CGTGCCTA | CTGCTGCCTT CCTTGGATG | 60 | – | – | 10 | Designed |
| *GAPDH* | NM_017008.4 | ACATCAAATGG GGTGATGCT | GTGGTTCACA CCCATCACAA | 62 | 10 | – | – | Designed |
| *GUSB* | NM_017015.3 | CATGACGAACC AGTCACCAC | ACGGTCTGCT TCCCATACAC | 64 | – | 50 | – | (Liu et al., 2020) |
| *YWAZ* | NM_013011.4 | TTGAGCAGAAG ACGGAAGGT | GAAGCATTGG GGATCAAGAA | 60 | 10 | – | 10 | Designed |
| *HSF1* | NM_024393.1 | GACATGAGCCT GCCTGACCT | GTTCAGTGCC CGTCAGAAGA | 60 | 150 | 100 | 100 | Designed |
| *DNAJB1* | NM_001395149.1 | TACCACCCGGA CAAGAACAAG | CAGGCCTTCC TCTCCATAGC | 64 | 50 | 50 | 50 | Designed |
| *HSPA1A* | NM_031971.2 | TGGTGCAGTCG GACATGAAG | GGTAGAACGA CCGGTTCTCG | 62 | 50 | 50 | 50 | Designed |
| *HSP90AA1* | S45392.1 | CTCGTCAAGAT GCCTGAGGAAG TGC | CTCCATGAAC GCCTTTGTAC CAGACTTAG | 64 | 10 | 10 | 10 | (Ammon-Treiber et al., 2004) |

**Table S3. Primer parameters for RT-qPCR in the rat model**. Targets are identified by gene name, NCBI accession number, and forward and reverse primer sequences. Product size (bp), forward and reverse melting temperatures (°C) and GC content (%), hairpin temperature (°C), and homodimer and heterodimer values (%) are listed. The sources of the primers (designed using NCBI or obtained from literature) are indicated. All primers target *Rattus norvegicus*.

| Gene | Product size (bp) | Melting temperature (°C) | GC content (%) | Hairpin temperature (°C) | Homodimer (%) | Heterodimer (%) |
|---|---|---|---|---|---|---|
| *18S rRNA* | 107 | 60.76 | 57.89 | 28.0 | 9.30 | 8.09 |
| *GAPDH* | 161 | 57.46 | 45.00 | 49.8 | 18.38 | 25.28 |
| *GUSB* | Unspecified | 62.60 | 55.00 | 48.0 | 14.96 | 11.63 |
| *YWAZ* | 136 | 58.95 | 50.00 | 18.7 | 9.50 | 14.23 |
| *HSF1* | 283 | 61.91 | 60.00 | 22.0 | 14.00 | 16.80 |
| *DNAJB1* | 162 | 59.93 | 63.13 | 30.0 | 23.60 | 27.50 |
| *HSPA1A* | 102 | 60.32 | 55.00 | 21.9 | 18.50 | 16.80 |
| *HSP90AA1* | Unspecified | 68.10 | 56.00 | 33.6 | 9.80 | 9.30 |

**Table S4. Western immunoblotting parameters for HSR targets in the rat model.** Targets are identified with the corresponding molecular weight (kDa), SDS-polyacrylamide gel percentage (%), amount of protein loaded (µg), SDS-polyacrylamide gel run time and voltage, transfer settings (time, voltage, amperage), casein blocking (percentage, time), primary antibody incubation parameters (concentration, time, temperature), and secondary antibody incubation parameters (concentration, time, temperature).

| Tissue | Target | Molecular weight (kDa) | Gel % | Protein loaded (µg) | SDS-polyacrylamide gel run | Transfer settings | Casein blocking | Primary antibody incubation | Secondary antibody incubation |
|---|---|---|---|---|---|---|---|---|---|
| Liver | HSF1 | 82 | 8 | 17.5 | 110 min, 180 V | 40 min, 25 V, 2.5 A | 1%, 30 min | 1:1000, 24h, 4 °C | 1:15,000, 45 min, 22 °C |
| | Hsp40 | 40 | 12 | 15 | 115 min, 180 V | 40 min, 25 V, 2.5 A | 20%, 1h | 1:1000, 24h, 4 °C | 1:15,000, 45 min, 22 °C |
| | Hsp70 | 72, 73 | 10 | 15 | 100 min, 180 V | 40 min, 25 V, 2.5 A | 1%, 30 min | 1:1000, 24h, 4 °C | 1:10,000, 45 min, 22 °C |
| | Hsp90 | 90 | 10 | 15 | 100 min, 180 V | 40 min, 25 V, 2.5 A | 1%, 30 min | 1:1000, 24h, 4 °C | 1:10,000, 45 min, 22 °C |
| Hypothalamus | HSF1 | 82 | 8 | 25 | 110 min, 180 V | 40 min, 25 V, 2.5 A | 2.5%, 30 min | 1:1000, 24h, 4 °C | 1:10,000, 1h, 22 °C |
| | Hsp40 | 40 | 15 | 25 | 110 min, 180 V | 40 min, 25 V, 2.5 A | 1%, 30 min | 1:1000, 1h, 22 °C | 1:10,000, 45 min, 22 °C |
| | Hsp70 | 72, 73 | 10 | 10 | 110 min, 180 V | 40 min, 25 V, 2.5 A | 1%, 30 min | 1:1000, 22h, 4 °C | 1:10,000, 45 min, 22 °C |
| | Hsp90 | 90 | 8 | 10 | 110 min, 180 V | 40 min, 25 V, 2.5 A | 1%, 30 min | 1:1000, 22h, 4 °C | 1:10,000, 45 min, 22 °C |
| Prefrontal cortex | HSF1 | 82 | 8 | 15 | 110 min, 180 V | 40 min, 25 V, 2.5 A | 1%, 30 min | 1:1000, 24h, 4 °C | 1:10,000, 45 min, 22 °C |
| | Hsp40 | 40 | 15 | 17.5 | 110 min, 180 V | 40 min, 25 V, 2.5 A | 1%, 30 min | 1:1000, 24h, 4 °C | 1:10,000, 45 min, 22 °C |
| | Hsp70 | 72, 73 | 10 | 15 | 110 min, 180 V | 40 min, 25 V, 2.5 A | 1%, 30 min | 1:1000, 24h, 4 °C | 1:10,000, 45 min, 22 °C |
| | Hsp90 | 90 | 8 | 15 | 110 min, 180 V | 40 min, 25 V, 2.5 A | 2.5%, 30 min | 1:1000, 24h, 4 °C | 1:10,000, 45 min, 22 °C |

**Table S5. Primary antibodies used for HSR protein targets in the rat model**. Information includes the commercial antibody name, protein target, purchasing company and catalogue number, antigen species targeted, molecular weight, clonality, and host isotype. All antibodies are cross-reactive and recognize *Homo sapiens* and *Rattus norvegicus*.

| Antibody name | Target | Company | Catalogue number | Antigen species | Molecular weight (kDa) | Clonality | Host Isotype |
|---|---|---|---|---|---|---|---|
| HSF1 Polyclonal Antibody | HSF1 | ProteinTech | #51034-1-AP | Human | 82 | Polyclonal | Rabbit IgG |
| HSP40 (C64B4) Rabbit mAb | Hsp40 | Cell Signalling | #4871 | Human | 40 | Monoclonal | Rabbit IgG |
| HSP70 Antibody | Hsp70 | Cell Signalling | #4872 | Human | 72, 73 | Polyclonal | Rabbit IgG |
| HSP90 (C45G5) Rabbit mAb | Hsp90 | Cell Signalling | #4877 | Human | 90 | Monoclonal | Rabbit IgG |

(i) Liver HSF1 ECL and Coomassie images

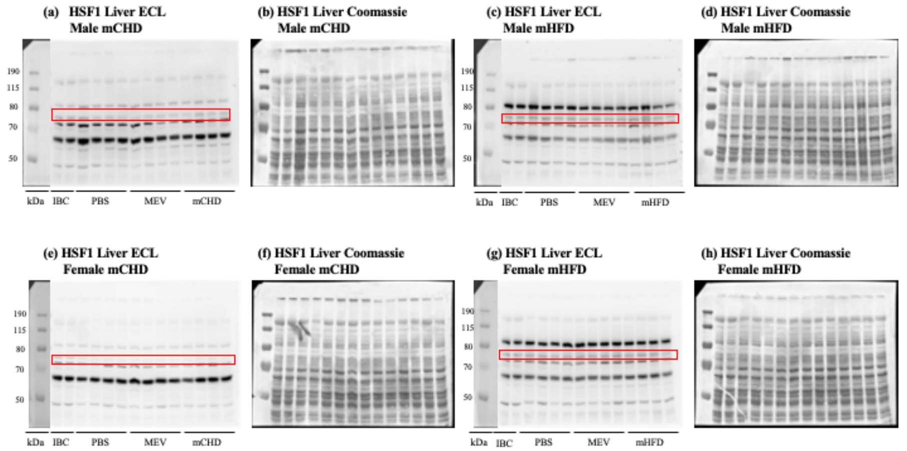


(ii) Hypothalamus HSF1 ECL and Coomassie images

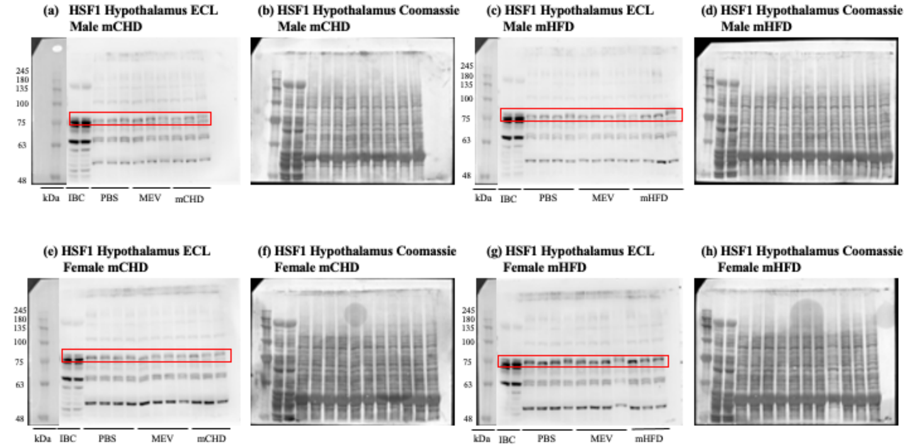


(iii) Prefrontal cortex HSF1 ECL and Coomassie images

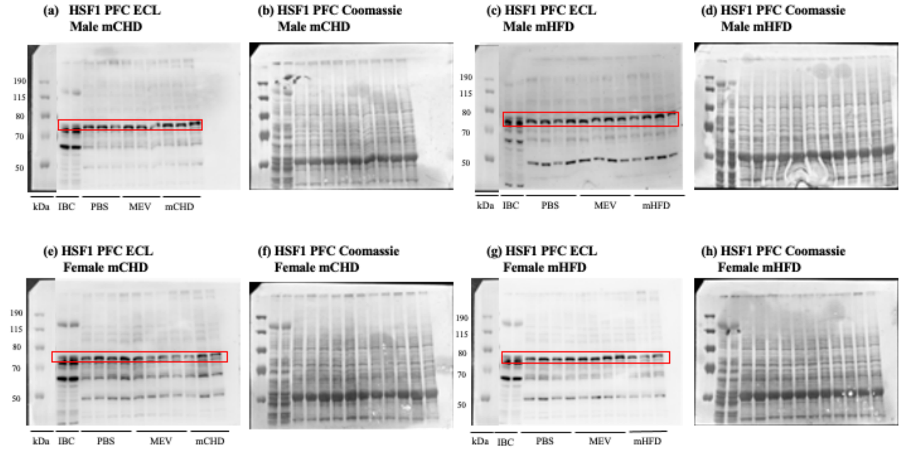


**Figure S4. ECL and Coomassie-stained immunoblot images of HSF1 in rat neonates**. (i) ECL and Coomassie-stained images for HSF1 in the liver. (ii) ECL and Coomassie-stained images for HSF1 in the hypothalamus. (iii) ECL and Coomassie-stained images for HSF1 in the prefrontal cortex.

**(i) Liver Hsp70 ECL and Coomassie images**

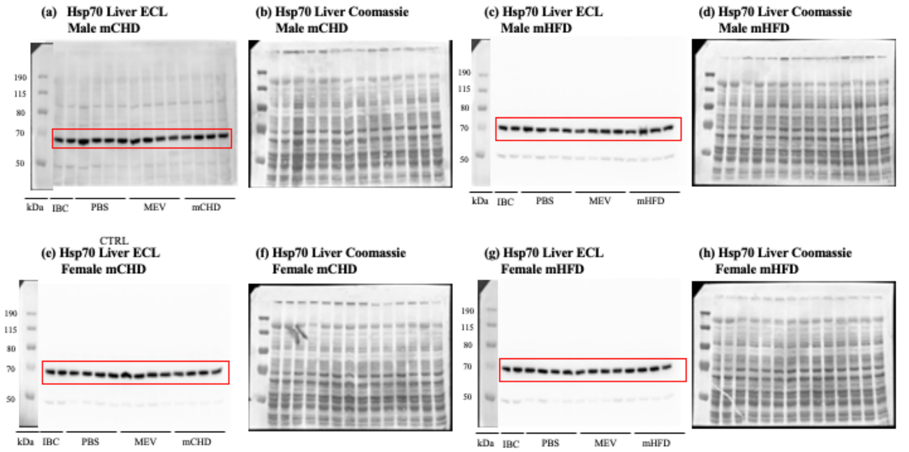


**(ii) Hypothalamus Hsp70 ECL and Coomassie images**

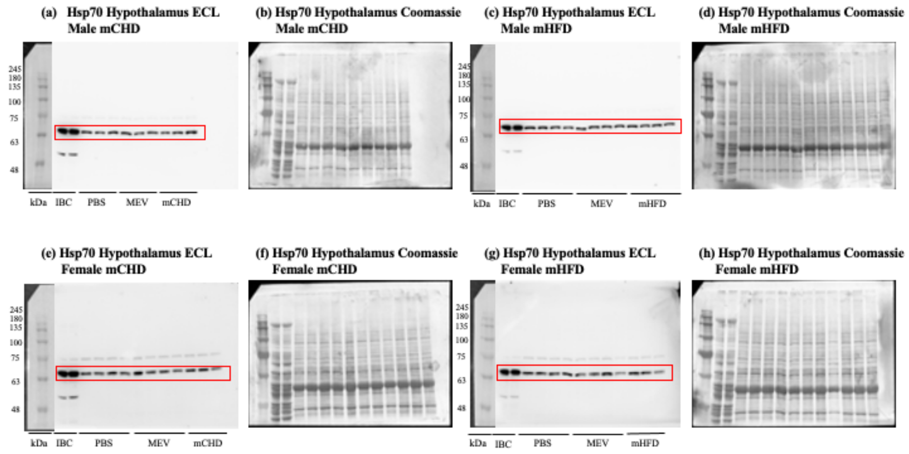


**(iii) Prefrontal cortex Hsp70 ECL and Coomassie images**

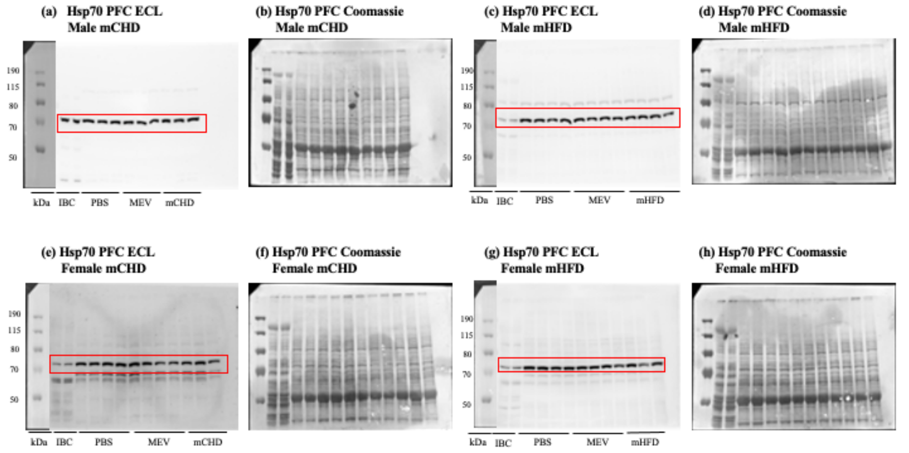


**Figure S5. ECL and Coomassie-stained immunoblot images of Hsp70 in rat neonates**. **(i)** ECL and Coomassie-stained images for Hsp70 in the liver. **(ii)** ECL and Coomassie-stained images for Hsp70 in the hypothalamus. **(iii)** ECL and Coomassie-stained images for Hsp70 in the prefrontal cortex.

(i) Liver Hsp90 ECL and Coomassie images

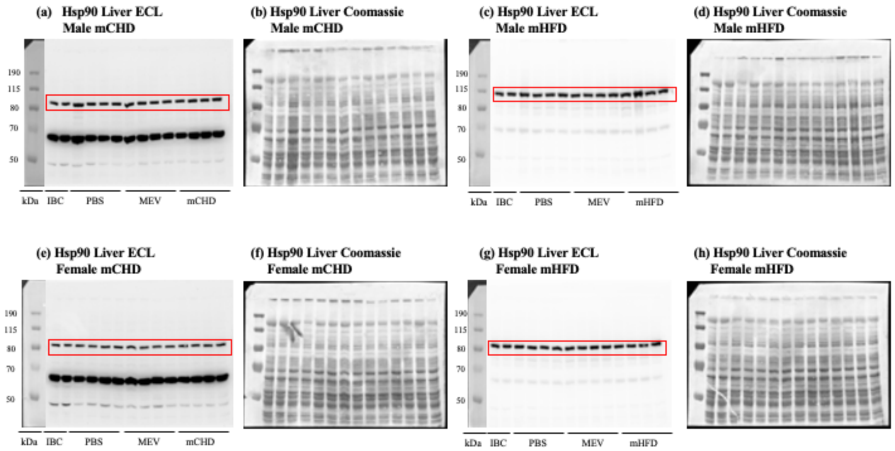


(ii) Hypothalamus Hsp90 ECL and Coomassie images

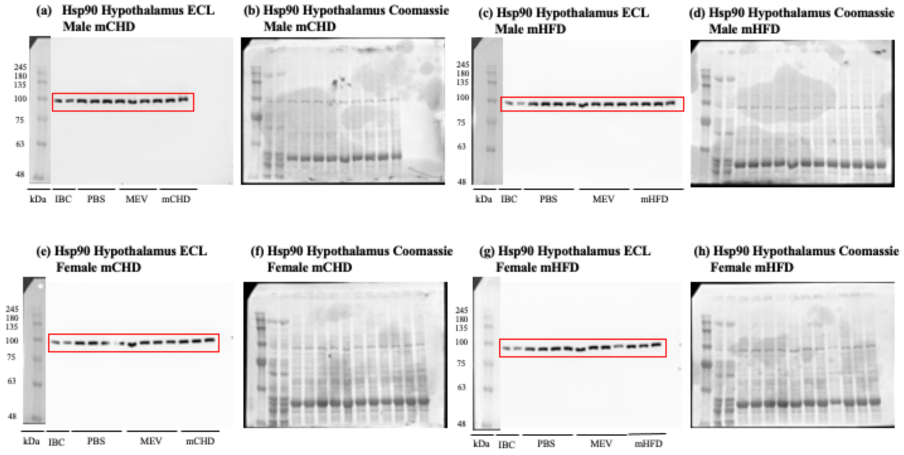


(iii) Prefrontal cortex Hsp90 ECL and Coomassie images

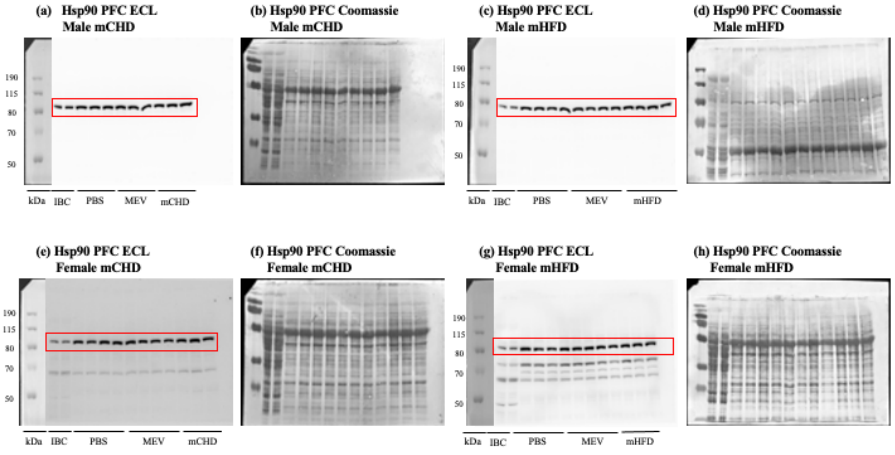


**Figure S6. ECL and Coomassie-stained immunoblot images of Hsp90 in rat neonates**. (i) ECL and Coomassie-stained images for Hsp90 in the liver. (ii) ECL and Coomassie-stained images for Hsp90 in the hypothalamus. (iii) ECL and Coomassie-stained images for Hsp90 in the prefrontal cortex.

(i) Liver Hsp40 ECL and Coomassie images

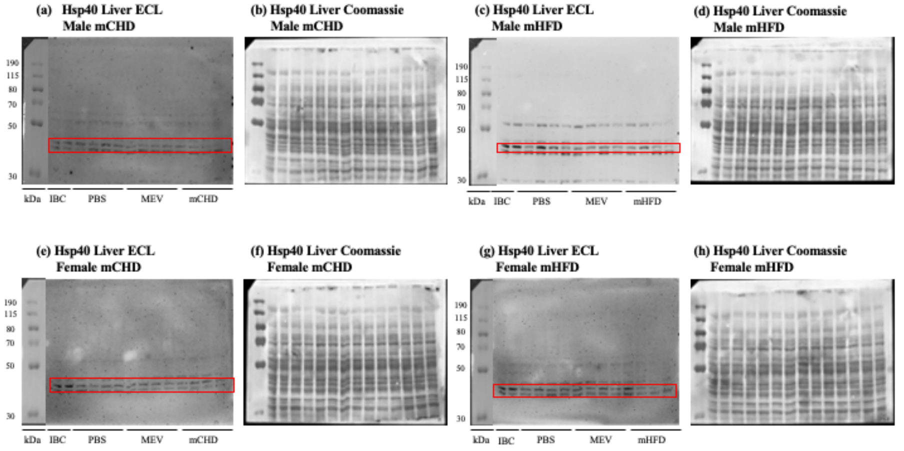


(ii) Hypothalamus Hsp40 ECL and Coomassie images

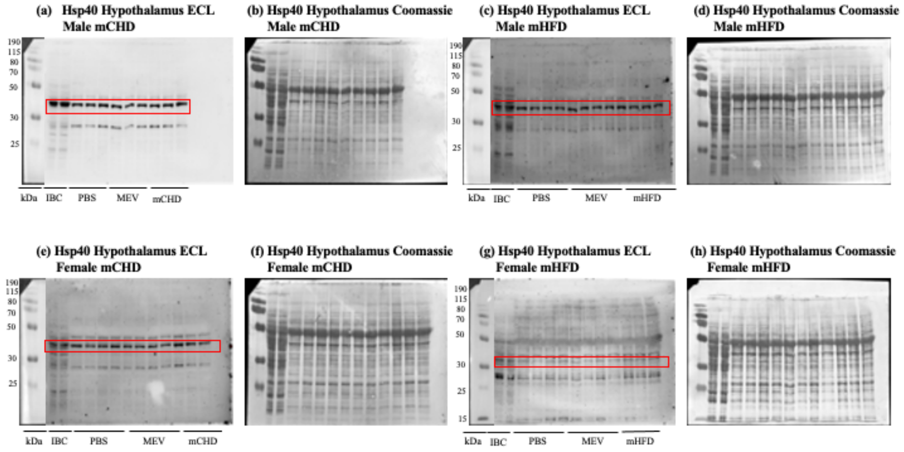


(iii) Prefrontal cortex Hsp40 ECL and Coomassie images

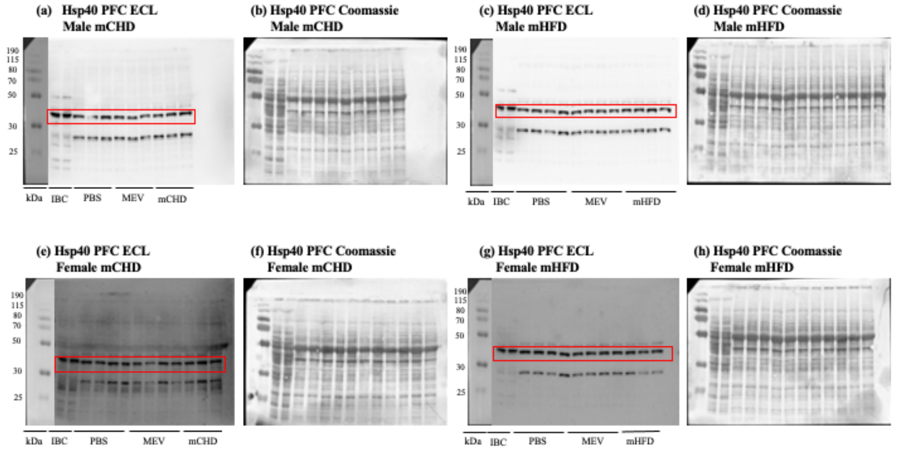


**Figure S7. ECL and Coomassie-stained immunoblot images of Hsp40 in rat neonates**. **(i)** ECL and Coomassie-stained images for Hsp40 in the liver. **(ii)** ECL and Coomassie-stained images for Hsp40 in the hypothalamus. **(iii)** ECL and Coomassie-stained images for Hsp40 in the prefrontal cortex.

**Table S6. Normalized transcript and protein data for HSR targets in the liver, hypothalamus and prefrontal cortex**. The data are organized by sex, treatment group, neonate ID, and tissue. Normalized transcript data are presented as quantity means. Normalized protein data are presented in atomic mass units.

| Sex | Treatment group | Neonate ID | Tissue | Normalized transcript (QM) | | | | Normalized protein (AU) | | | |
|---|---|---|---|---|---|---|---|---|---|---|---|
| | | | | HSF1 | HSPA1A | HSP90AA1 | DNAJB1 | HSF1 | Hsp70 | Hsp90 | Hsp40 |
| **Male** | mCHD-PBS | D1M1 | Liver | 2.436188848 | 1.656769374 | 1.449968241 | 3.009002977 | 0.171898617 | 0.099025771 | 0.387089964 | 6.749707468 |
| **Male** | mCHD-PBS | D3M1 | Liver | 5.419481327 | 2.986506676 | 1.288870824 | 7.399306052 | 0.089537186 | 0.192480401 | 0.038332484 | 6.797471077 |
| **Male** | mCHD-PBS | D9M1 | Liver | 4.34711594 | 4.324865352 | 1.198930151 | 11.82736006 | 0.014259119 | 0.280109431 | 0.35972159 | 6.840995982 |
| **Male** | mCHD-PBS | D11M2 | Liver | 3.430277295 | 1.721399638 | 1.09695633 | 7.249028719 | 0.039679112 | 0.425311417 | 0.964569341 | 6.946522949 |
| **Male** | mCHD-MEV | D1M4 | Liver | 4.822682603 | 2.475993105 | 0.823033445 | 5.694322305 | 0.062414331 | 0.283261389 | 0.098926056 | 7.079112699 |
| **Male** | mCHD-MEV | D3M2 | Liver | 6.11217303 | 2.57986536 | 1.723115954 | 10.45421477 | 0.105931269 | 0.252280855 | 0.275059214 | 7.040587826 |
| **Male** | mCHD-MEV | D9M4 | Liver | 1.647635467 | 1.792131561 | 1.310328017 | 3.779316945 | 0.126325237 | 0.036824978 | 0.262015768 | 7.033418042 |
| **Male** | mCHD-MEV | D11M3 | Liver | 5.200892948 | 2.309472209 | 1.266695013 | 10.10532823 | 0.133369688 | 0.05820046 | 0.382079166 | 6.993794465 |
| **Male** | mCHD | D1M6 | Liver | 5.367685157 | 2.107326421 | 1.315803555 | 8.57526999 | 0.090448443 | 0.103886142 | 0.470055372 | 7.041895073 |
| **Male** | mCHD | D3M3 | Liver | 2.149643225 | 1.406666738 | 0.551054958 | 5.317073021 | 0.057783341 | 0.059347201 | 0.219806819 | 7.081977214 |
| **Male** | mCHD | D9M6 | Liver | 2.03058354 | 1.033900936 | 1.207235253 | 6.242194106 | 0.101006959 | 0.088088643 | 0.159686289 | 6.896427317 |
| **Male** | mCHD | D11M6 | Liver | 6.959875935 | 3.452134788 | 0.656432048 | 12.64699561 | 0.107326728 | 0.053635318 | 0.274345 | 6.704247853 |
| **Male** | mHFD-PBS | D2M2 | Liver | 2.321029164 | 0.768827891 | 1.147894286 | 1.831698353 | 0.297908267 | 0.346392937 | 0.407000924 | 6.009843098 |
| **Male** | mHFD-PBS | D4M1 | Liver | 3.988232555 | 2.223408606 | 0.8546777 | 5.597123831 | 0.138944986 | 0.105046756 | 1.046983609 | 5.868583568 |
| **Male** | mHFD-PBS | D6M1 | Liver | 10.404798 | 5.024238206 | 1.096068025 | 10.18520886 | 0.13954249 | 0.140143654 | 1.066341693 | 6.028100172 |
| **Male** | mHFD-PBS | D8M1 | Liver | 6.279454336 | 2.990447947 | 0.846671347 | 9.904507254 | 0.110068515 | 0.153923177 | 0.432133064 | 6.076395035 |
| **Male** | mHFD-MEV | D2M4 | Liver | 2.356714821 | 0.982701532 | 0.91826876 | 2.533495959 | 0.000678528 | 0.165410512 | 0.405813907 | 6.317859106 |
| **Male** | mHFD-MEV | D4M3 | Liver | 7.656742884 | 2.978997639 | 1.012834315 | 7.637730208 | 0.059335949 | 0.091202206 | 0.187306497 | 6.588806964 |
| **Male** | mHFD-MEV | D6M3 | Liver | 32.53059953 | 13.19449562 | 0.777728107 | 18.38394664 | 0.027635266 | 0.01590096 | 0.331540686 | 6.068797562 |
| **Male** | mHFD-MEV | D8M4 | Liver | 3.937226713 | 1.291570396 | 1.246961094 | 3.082840521 | 0.051171248 | 0.094932374 | 0.24927669 | 6.410268176 |
| **Male** | mHFD | D2M7 | Liver | 6.121801913 | 1.883754221 | 0.778581352 | 3.631151233 | 0.211102749 | 0.245249262 | 0.049673666 | 6.408172676 |
| **Male** | mHFD | D4M6 | Liver | 12.81249362 | 3.780972705 | 1.010722403 | 6.989209362 | 0.172805929 | 0.085415114 | 0.093607747 | 6.366097572 |
| **Male** | mHFD | D6M5 | Liver | 11.18804511 | 4.260548148 | 0.852294934 | 19.94826921 | 0.110813666 | 0.023969282 | 0.500117093 | 5.871464351 |
| **Male** | mHFD | D8M5 | Liver | 3.408277122 | 1.810829048 | 0.590464957 | 1.940861918 | 0.011093143 | 0.046633958 | 0.484871988 | 6.107678737 |
| **Female** | mCHD-PBS | D1F1 | Liver | 6.509011829 | 1.983938931 | 1.023725409 | 3.918483505 | 0.416438963 | 0.355440919 | 1.258467115 | 6.539005691 |
| **Female** | mCHD-PBS | D3F1 | Liver | 6.201574977 | 2.361234312 | 1.641416068 | 22.15724962 | 0.879314162 | 1.517737034 | 0.496052751 | 6.802529321 |
| **Female** | mCHD-PBS | D9F1 | Liver | 2.999084498 | 24.21721077 | 1.101668657 | 4.315399801 | 0.653735553 | 1.315957389 | 0.663797144 | 6.865609282 |
| **Female** | mCHD-PBS | D11F2 | Liver | 3.478318138 | 3.327938578 | 0.711366877 | 9.41942024 | 0.352685705 | 1.170031241 | 0.76281329 | 6.994238948 |
| **Female** | mCHD-MEV | D1F3 | Liver | 6.51297937 | 3.031775231 | 1.242331552 | 5.555921951 | 0.338255613 | 1.545880693 | 0.716853207 | 7.081918703 |
| **Female** | mCHD-MEV | D3F5 | Liver | 8.732118861 | 3.747232732 | 1.447410539 | 10.3275388 | 0.366319067 | 1.362044795 | 0.762452776 | 7.000208681 |
| **Female** | mCHD-MEV | D9F3 | Liver | 6.260668166 | 2.968268105 | 1.003402785 | 6.803817784 | 0.678347908 | 2.215182102 | 0.30376846 | 7.034806268 |
| **Female** | mCHD-MEV | D11F4 | Liver | 4.826681238 | 2.180251953 | 1.177152993 | 5.853381766 | 0.272821504 | 1.40333353 | 0.493267744 | 7.139779002 |
| **Female** | mCHD | D1F6 | Liver | 2.299301111 | 0.977249466 | 1.064059481 | 2.221513437 | 0.458777327 | 0.882817377 | 0.650035912 | 7.105908278 |

| | | | | | | | | | | | |
|---|---|---|---|---|---|---|---|---|---|---|---|
| **Female** | mCHD | D3F7 | Liver | 1.0764101 | 1.092129971 | 1.048313936 | 2.571861009 | 1.012757568 | 1.321771141 | 0.519493531 | 6.965245341 |
| **Female** | mCHD | D9F5 | Liver | 0.628498764 | 0.583772669 | 0.932449412 | 1.690765133 | 1.037031513 | 0.968615308 | 0.65581012 | 6.997623382 |
| **Female** | mCHD | D11F6 | Liver | 2.169343743 | 0.639403664 | 1.361314981 | 1.525251315 | 1.272259351 | 1.633533231 | 0.45373308 | 6.841886542 |
| **Female** | mHFD-PBS | D2F2 | Liver | 3.883521859 | 1.324523973 | 0.830165672 | 4.766178003 | 1.000932783 | 1.986597672 | 0.749759496 | 8.980769257 |
| **Female** | mHFD-PBS | D4F1 | Liver | 3.364188371 | 1.621971923 | 1.378457888 | 3.748512016 | 1.287136182 | 2.419520339 | 1.214028538 | 12.50876926 |
| **Female** | mHFD-PBS | D6F1 | Liver | 2.043605315 | 1.169960193 | 0.84886723 | 2.976542347 | 1.336288898 | 2.410285713 | 1.573228879 | 20.93476926 |
| **Female** | mHFD-PBS | D8F2 | Liver | 2.144952128 | 1.463122874 | 1.132381653 | 1.880008445 | 0.77434137 | 1.17872683 | 0.113798918 | 22.15376926 |
| **Female** | mHFD-MEV | D2F3 | Liver | 4.658570605 | 2.362542451 | 0.771799543 | 3.179701493 | 1.370269524 | 1.503167654 | 0.344609615 | 23.13476926 |
| **Female** | mHFD-MEV | D4F4 | Liver | 1.710128066 | 2.372496655 | 0.577886864 | 4.144546866 | 0.862300547 | 1.23582232 | 0.270749268 | 16.02376926 |
| **Female** | mHFD-MEV | D6F4 | Liver | 1.859877938 | 0.78679627 | 0.956371764 | 1.720851699 | 0.675992338 | 0.718306858 | 0.061049874 | 17.29876926 |
| **Female** | mHFD-MEV | D8F4 | Liver | 2.794573278 | 0.775271005 | 1.122486987 | 1.960058799 | 0.928689588 | 0.8019335 | 0.471998335 | 12.26876926 |
| **Female** | mHFD | D2F5 | Liver | 2.68427817 | 0.820458221 | 1.298941977 | 1.799464352 | 0.838355372 | 1.235534152 | 0.077451002 | 36.34476926 |
| **Female** | mHFD | D4F6 | Liver | 8.818271873 | 2.303645126 | 1.329132891 | 4.752590304 | 1.171614271 | 1.833844926 | 0.479272738 | 11.65976926 |
| **Female** | mHFD | D6F6 | Liver | 5.259360153 | 1.0915065 | 1.087416615 | 3.47937366 | 1.135969776 | 1.917227721 | 0.352681963 | 8.396769257 |
| **Female** | mHFD | D8F5 | Liver | 5.29882552 | 1.094412117 | 1.176098323 | 1.849397476 | 0.400903744 | 0.550394485 | 0.303748425 | 17.97176926 |
| **Male** | mCHD-PBS | D1M1 | Hypothalamus | 0.968544788 | 0.892643833 | 0.149969381 | 1.977430931 | 0.223826692 | 0.227368105 | 1.22301346 | 5.658601911 |
| **Male** | mCHD-PBS | D3M1 | Hypothalamus | 0.610806156 | 0.727525733 | 0.09352309 | 1.427472808 | 0.239699653 | 0.538809845 | 1.11539194 | 6.061176888 |
| **Male** | mCHD-PBS | D9M1 | Hypothalamus | 0.304352082 | 0.764168623 | 0.130444446 | 1.453661216 | 0.253228446 | 0.46567697 | 1.39282364 | 1.348707227 |
| **Male** | mCHD-PBS | D11M2 | Hypothalamus | 0.454699267 | 0.545858081 | 0.143224834 | 1.456744761 | | | | |
| **Male** | mCHD-MEV | D1M4 | Hypothalamus | 0.729588595 | 0.894343595 | 0.108125787 | 2.049380953 | 0.411936043 | 0.458166063 | 1.40618722 | 2.184713019 |
| **Male** | mCHD-MEV | D3M2 | Hypothalamus | 0.914619917 | 1.007248823 | 0.128349697 | 3.094863951 | 0.259797218 | 0.8959676 | 1.43954939 | 2.350554405 |
| **Male** | mCHD-MEV | D9M4 | Hypothalamus | 0.578660448 | 0.926077954 | | 2.133835409 | 0.080929588 | 0.842484346 | 1.60084426 | 2.38208313 |
| **Male** | mCHD-MEV | D11M3 | Hypothalamus | 0.592704363 | 0.758015916 | 0.103161987 | 2.038354847 | | | | |
| **Male** | mCHD | D1M6 | Hypothalamus | 0.856901851 | 0.786599663 | 0.134144071 | 2.114363265 | 0.108588639 | 0.816760093 | 2.28444196 | 3.466905587 |
| **Male** | mCHD | D3M3 | Hypothalamus | 0.519977094 | 0.738843907 | 0.108443053 | 2.121926357 | 0.286188898 | 0.6323315 | 2.31012555 | 2.340201399 |
| **Male** | mCHD | D9M6 | Hypothalamus | 0.3786474 | 0.673924872 | 0.104077886 | 2.186072166 | 0.063179134 | 0.254871873 | 2.17346744 | 2.168785239 |
| **Male** | mCHD | D11M6 | Hypothalamus | 0.628615139 | 0.794681043 | 0.077278033 | 2.077011128 | | | | |
| **Male** | mHFD-PBS | D2M2 | Hypothalamus | 1.882889217 | 5.317512591 | 2.244890837 | 2.264780247 | 0.002303693 | 0.464799404 | 1.10866654 | 1.534280168 |
| **Male** | mHFD-PBS | D4M1 | Hypothalamus | 0.724317065 | 0.868116609 | 0.210238396 | 2.772654305 | 0.014669528 | 0.622499667 | 1.32707239 | 0.565826648 |
| **Male** | mHFD-PBS | D6M1 | Hypothalamus | 0.707838296 | 0.650410395 | 0.182646085 | 2.190033625 | 0.033908088 | 0.724655706 | 1.48810128 | 0.794383647 |
| **Male** | mHFD-PBS | D8M1 | Hypothalamus | 0.627596963 | 0.725575015 | 0.166609849 | 1.970264947 | 0.05382539 | 0.915099941 | 1.33108789 | 1.749287879 |
| **Male** | mHFD-MEV | D2M4 | Hypothalamus | 0.756350896 | 0.555932307 | 0.22643342 | 2.055103527 | 0.111499402 | 0.843843211 | 1.65566019 | 0.975094028 |
| **Male** | mHFD-MEV | D4M3 | Hypothalamus | 0.752767925 | 0.674677285 | 0.244395108 | 2.450702006 | 0.098623927 | 0.517819896 | 1.34899424 | 0.947329516 |
| **Male** | mHFD-MEV | D6M3 | Hypothalamus | 0.724220813 | 0.681289215 | 0.220501446 | 2.155941592 | 0.150554466 | 0.461969844 | 2.72724853 | 1.042412752 |
| **Male** | mHFD-MEV | D8M4 | Hypothalamus | 0.76856105 | 0.820302889 | 0.201947531 | 1.70101913 | 0.153784789 | 0.409777606 | 2.68796695 | 0.902592582 |
| **Male** | mHFD | D2M7 | Hypothalamus | 0.745504357 | 0.688821246 | 0.186305504 | 1.739724989 | 0.25869241 | 0.071931246 | 2.24898088 | 1.413658981 |
| **Male** | mHFD | D4M6 | Hypothalamus | 0.757335379 | 0.643354193 | 0.120399962 | 1.41433674 | 0.057196624 | 0.262671786 | 2.87819899 | 1.425182066 |
| **Male** | mHFD | D6M5 | Hypothalamus | 0.899356759 | 0.821949258 | 0.201260676 | 1.748489391 | 0.124758471 | 0.055334007 | 2.82586867 | 0.454504606 |

| | | | | | | | | | | | |
|---|---|---|---|---|---|---|---|---|---|---|---|
| **Male** | mHFD | D8M5 | Hypothalamus | 0.855993848 | 0.794010307 | 0.109558843 | 1.865553577 | 0.083677825 | 0.550630024 | 1.90071928 | 1.707828902 |
| **Female** | mCHD-PBS | D1F1 | Hypothalamus | 0.980524948 | 0.583266827 | 0.115886975 | 1.404030363 | 0.089224006 | 0.641195484 | 3.150537397 | 0.522018055 |
| **Female** | mCHD-PBS | D3F1 | Hypothalamus | 0.698349528 | 0.910272144 | 0.100151858 | 1.629360948 | 0.028503721 | 0.821445445 | 3.337705476 | 0.084177719 |
| **Female** | mCHD-PBS | D9F1 | Hypothalamus | 0.400904684 | 0.853868583 | 0.121284004 | 1.523255387 | 0.033192272 | 0.819022368 | 0.689062145 | 0.273966867 |
| **Female** | mCHD-PBS | D11F2 | Hypothalamus | 0.575863438 | 0.764547888 | 0.129451243 | 1.467920183 | 0.080677582 | 0.880857936 | 0.11610317 | 0.387474715 |
| **Female** | mCHD-MEV | D1F3 | Hypothalamus | 0.632203942 | 0.57737302 | 0.126134353 | 1.515037372 | 0.102042023 | 0.828597753 | 1.193279737 | 0.320854041 |
| **Female** | mCHD-MEV | D3F5 | Hypothalamus | 0.652452999 | 1.052693218 | 0.134046261 | 1.606953369 | 0.184468179 | 0.822877734 | 2.516206885 | 0.269842574 |
| **Female** | mCHD-MEV | D9F3 | Hypothalamus | 0.684147398 | 1.078988484 | 0.121180318 | 1.958005181 | 0.188589119 | 0.866486799 | 2.893439268 | 0.072859706 |
| **Female** | mCHD-MEV | D11F4 | Hypothalamus | 0.699177986 | 0.75235774 | 0.144926449 | 1.67563047 | 0.184788 | 0.91265854 | 2.713816489 | 0.085136494 |
| **Female** | mCHD | D1F6 | Hypothalamus | 0.584910339 | 0.784557148 | 0.09066147 | 1.787280386 | 0.09256755 | 0.864557856 | 2.152427146 | 0.248560366 |
| **Female** | mCHD | D3F7 | Hypothalamus | 0.461107561 | 1.020294239 | 0.145044741 | 1.876361881 | 0.184236716 | 0.833655174 | 1.409742388 | 0.240457367 |
| **Female** | mCHD | D9F5 | Hypothalamus | 0.436480312 | 0.825601875 | 0.158890848 | 1.646941047 | 0.081040905 | 1.025743992 | 1.721034553 | 0.051908383 |
| **Female** | mCHD | D11F6 | Hypothalamus | 0.976217711 | 0.853319711 | 0.15465158 | 1.714251811 | | | | |
| **Female** | mHFD-PBS | D2F2 | Hypothalamus | 0.946148608 | 0.855077005 | 0.181787765 | 1.571838706 | 0.389944712 | 0.314133613 | 1.599075292 | 0.427993898 |
| **Female** | mHFD-PBS | D4F1 | Hypothalamus | 0.660066853 | 0.856590366 | 0.345446236 | 1.987349127 | 0.213846052 | 0.4544623 | 1.787113007 | 0.410537928 |
| **Female** | mHFD-PBS | D6F1 | Hypothalamus | 0.903334507 | 1.103604287 | 0.205623863 | 2.310879083 | 0.302273926 | 0.59531971 | 1.65433247 | 0.369529291 |
| **Female** | mHFD-PBS | D8F2 | Hypothalamus | 0.633048454 | 0.884632072 | 0.182872205 | 1.555590554 | 0.237746395 | 0.614153572 | 1.132125479 | 0.34514778 |
| **Female** | mHFD-MEV | D2F3 | Hypothalamus | 0.839423639 | 1.187341763 | 0.328478429 | 2.257858824 | 0.254636276 | 0.555394106 | 2.139869112 | 0.342351993 |
| **Female** | mHFD-MEV | D4F4 | Hypothalamus | 0.713374934 | 1.3226256 | 0.226434375 | 2.581340016 | 0.068644624 | 0.475760306 | 1.865111243 | 0.379854226 |
| **Female** | mHFD-MEV | D6F4 | Hypothalamus | 0.810644513 | 0.931506849 | 0.297845211 | 1.65382142 | 0.265480537 | 0.285305004 | 1.729275271 | 0.36853522 |
| **Female** | mHFD-MEV | D8F4 | Hypothalamus | 0.781012101 | 0.988340845 | 0.207062695 | 1.68725534 | 0.346208367 | 0.181507572 | 2.195569341 | 0.308106766 |
| **Female** | mHFD | D2F5 | Hypothalamus | 0.773579724 | 0.852930184 | 0.177097822 | 1.885571851 | 0.437128544 | 0.133730701 | 1.537622051 | 0.224840885 |
| **Female** | mHFD | D4F6 | Hypothalamus | 0.86345216 | 1.129018994 | 0.166319772 | 1.765911404 | 0.246507834 | 0.51379228 | 1.332266316 | 0.290761488 |
| **Female** | mHFD | D6F6 | Hypothalamus | 0.993920802 | 0.807581551 | 0.264399222 | 1.246270469 | 0.350556839 | 0.809842552 | 1.772890731 | 0.180666796 |
| **Female** | mHFD | D8F5 | Hypothalamus | 0.988796238 | 1.013964103 | 0.246603735 | 1.255331962 | | | | |
| **Male** | mCHD-PBS | D1M1 | Prefrontal cortex | 8.368176262 | 8.846225289 | 2.95346643 | 10.80546988 | 1.273658557 | 0.211265881 | 0.72358355 | 0.534431909 |
| **Male** | mCHD-PBS | D3M1 | Prefrontal cortex | 7.570211586 | 9.283857482 | 1.845430561 | 11.56033075 | 1.15321753 | 0.472590641 | 0.950440592 | 0.664175353 |
| **Male** | mCHD-PBS | D9M1 | Prefrontal cortex | 4.049381555 | 13.22000696 | 1.875573167 | 10.60815791 | 0.186779554 | 0.46876523 | 0.67094412 | 0.044402197 |
| **Male** | mCHD-PBS | D11M2 | Prefrontal cortex | 4.205277286 | 8.634104047 | 1.408632191 | 11.12823664 | | | | |
| **Male** | mCHD-MEV | D1M4 | Prefrontal cortex | 8.563206475 | 9.196419743 | 1.745055597 | 10.73988614 | 0.453653914 | 0.340718672 | 0.517572064 | 0.443341659 |
| **Male** | mCHD-MEV | D3M2 | Prefrontal cortex | 7.205176899 | 9.496809833 | 1.42095352 | 13.92975254 | 0.2955689 | 0.357053875 | 0.407027248 | 0.508964742 |
| **Male** | mCHD-MEV | D9M4 | Prefrontal cortex | 7.253955046 | 11.35698811 | 1.731134779 | 12.38033603 | 0.009216253 | 0.53132906 | 0.354899332 | 0.282462545 |
| **Male** | mCHD-MEV | D11M3 | Prefrontal cortex | 8.170499469 | 11.11519554 | 1.533452797 | 13.98804925 | | | | |
| **Male** | mCHD | D1M6 | Prefrontal cortex | 7.536446523 | 7.094845765 | 1.616663749 | 11.33191577 | 0.557506027 | 0.712756785 | 0.451653613 | 0.225151439 |
| **Male** | mCHD | D3M3 | Prefrontal cortex | 4.041479729 | 7.929480973 | 1.429478182 | 12.96804103 | 0.491908886 | 0.530830093 | 0.610642839 | 1.1321965 |
| **Male** | mCHD | D9M6 | Prefrontal cortex | 3.770923308 | 7.903527908 | 1.337358591 | 12.20796276 | 0.966914811 | 0.62106737 | 0.879892235 | 1.855468424 |
| **Male** | mCHD | D11M6 | Prefrontal cortex | 9.543417227 | 10.98398794 | 1.827727424 | 16.03799502 | | | | |
| **Male** | mHFD-PBS | D2M2 | Prefrontal cortex | 6.891979197 | 7.945740278 | 1.578893366 | 11.85255238 | 0.643915909 | 1.981429508 | 1.346986041 | 2.581175442 |

| | | | | | | | | | | | |
|---|---|---|---|---|---|---|---|---|---|---|---|
| **Male** | mHFD-PBS | D4M1 | Prefrontal cortex | 6.560555539 | 9.319541165 | 1.554715055 | 12.22042888 | 0.478989265 | 0.70391638 | 0.673306522 | 2.288060732 |
| **Male** | mHFD-PBS | D6M1 | Prefrontal cortex | 8.639063297 | 9.862089149 | 1.54103033 | 12.63112764 | 0.355748375 | 0.430117995 | 0.498844463 | 3.636074151 |
| **Male** | mHFD-PBS | D8M1 | Prefrontal cortex | 8.906330474 | 9.871532851 | 1.675198168 | 14.18958727 | 0.446181683 | 0.487516178 | 0.550866716 | 3.382706219 |
| **Male** | mHFD-MEV | D2M4 | Prefrontal cortex | 8.96440191 | 8.806135414 | 1.735645039 | 14.54118582 | 0.211069922 | 0.387355141 | 0.434062143 | 1.287609974 |
| **Male** | mHFD-MEV | D4M3 | Prefrontal cortex | 5.900264552 | 7.812287237 | 1.390200967 | 10.89059529 | 0.324711026 | 0.312424775 | 0.356096494 | 2.000164387 |
| **Male** | mHFD-MEV | D6M3 | Prefrontal cortex | 9.125081998 | 9.569734949 | 1.647422192 | 14.09895429 | 0.470408888 | 0.419183117 | 0.426623672 | 1.827716267 |
| **Male** | mHFD-MEV | D8M4 | Prefrontal cortex | 9.485542543 | 11.35978993 | 2.017579561 | 12.07155246 | 0.266549067 | 0.361642787 | 0.435639972 | 1.523420244 |
| **Male** | mHFD | D2M7 | Prefrontal cortex | 7.061169939 | 8.055473786 | 1.384492704 | 12.35585659 | 0.154617919 | 0.511614249 | 0.592985497 | 2.784387881 |
| **Male** | mHFD | D4M6 | Prefrontal cortex | 8.196095863 | 8.806163256 | 1.83941608 | 10.68182199 | 0.57029266 | 0.543649073 | 0.58504273 | 2.74620152 |
| **Male** | mHFD | D6M5 | Prefrontal cortex | 10.41961919 | 8.424183151 | 1.768626689 | 8.989444004 | 0.625300012 | 0.566304992 | 0.831139794 | 2.826582371 |
| **Male** | mHFD | D8M5 | Prefrontal cortex | 9.190293332 | 8.412708258 | 1.578445046 | 8.444863103 | 0.443508975 | 0.773167296 | 1.37916322 | 3.056167215 |
| **Female** | mCHD-PBS | D1F1 | Prefrontal cortex | 11.74370892 | 8.547801774 | 1.692523685 | 13.95961603 | 0.205849769 | 0.602917721 | 1.706016408 | 0.524757865 |
| **Female** | mCHD-PBS | D3F1 | Prefrontal cortex | 14.84053857 | 14.53452121 | 2.651692531 | 16.37752616 | 0.442083077 | 0.671280393 | 1.850397947 | 0.016194656 |
| **Female** | mCHD-PBS | D9F1 | Prefrontal cortex | 6.986431493 | 12.78551994 | 1.900689931 | 14.00622912 | 0.187967883 | 0.503127767 | 1.596642543 | 0.276085331 |
| **Female** | mCHD-PBS | D11F2 | Prefrontal cortex | 7.829372369 | 13.77195281 | 1.868957807 | 12.61591038 | 0.393834871 | 0.65695141 | 1.540755689 | 0.628797009 |
| **Female** | mCHD-MEV | D1F3 | Prefrontal cortex | 9.964852749 | 11.90806036 | 1.930791869 | 17.18570984 | 0.101661401 | 1.124473723 | 1.916971169 | 0.658498255 |
| **Female** | mCHD-MEV | D3F5 | Prefrontal cortex | 11.2642518 | 14.35143764 | 1.981646294 | 16.20430909 | 0.17642305 | 0.771733788 | 1.748831537 | 0.706792155 |
| **Female** | mCHD-MEV | D9F3 | Prefrontal cortex | 10.69212933 | 11.53900944 | 2.304740418 | 14.0743143 | 0.086783811 | 1.282238731 | 1.612238785 | 0.225636297 |
| **Female** | mCHD-MEV | D11F4 | Prefrontal cortex | 7.89789602 | 9.437861022 | 2.191292971 | 7.804321364 | 0.104035534 | 1.052662192 | 2.916733224 | 0.577404168 |
| **Female** | mCHD | D1F6 | Prefrontal cortex | 11.64641037 | 10.82893025 | 2.235874694 | 16.04515493 | 0.221569049 | 0.912582001 | 2.545183888 | 0.552442674 |
| **Female** | mCHD | D3F7 | Prefrontal cortex | 7.402210431 | 13.00090198 | 2.239529591 | 18.38881227 | 0.392310567 | 0.934512718 | 2.206064214 | 0.385009081 |
| **Female** | mCHD | D9F5 | Prefrontal cortex | 5.989594257 | 12.28611699 | 1.781469397 | 12.38376384 | 0.593633093 | 0.839795494 | 2.747638794 | 1.222650872 |
| **Female** | mCHD | D11F6 | Prefrontal cortex | 17.9395937 | 22.60902455 | 2.378834372 | 20.80090637 | | | | |
| **Female** | mHFD-PBS | D2F2 | Prefrontal cortex | 9.230882487 | 11.32464139 | 2.129606693 | 15.18227323 | 0.392680874 | 0.879315547 | 2.016985143 | 1.926454349 |
| **Female** | mHFD-PBS | D4F1 | Prefrontal cortex | 7.81768934 | 12.98854828 | 2.241963126 | 15.38747421 | 0.514014883 | 0.621791053 | 1.230521337 | 1.834129479 |
| **Female** | mHFD-PBS | D6F1 | Prefrontal cortex | 11.16040058 | 11.9380012 | 2.228170728 | 13.71230284 | 0.309024537 | 0.718131541 | 1.482608195 | 1.95766162 |
| **Female** | mHFD-PBS | D8F2 | Prefrontal cortex | 9.770433411 | 11.40665937 | 2.163102614 | 12.10151434 | 0.487643903 | 0.759612621 | 1.776763479 | 1.360720183 |
| **Female** | mHFD-MEV | D2F3 | Prefrontal cortex | 7.837300743 | 9.761724963 | 1.846059211 | 11.70719963 | 0.489311079 | 0.839245916 | 1.782125458 | 1.058230056 |
| **Female** | mHFD-MEV | D4F4 | Prefrontal cortex | 8.410559561 | 13.77108694 | 2.250025734 | 16.08506202 | 0.374319862 | 1.257704839 | 2.429727748 | 0.766620414 |
| **Female** | mHFD-MEV | D6F4 | Prefrontal cortex | 10.79953439 | 11.48281929 | 2.097562823 | 14.14181727 | 0.593879497 | 1.054439767 | 2.126697786 | 0.714155333 |
| **Female** | mHFD-MEV | D8F4 | Prefrontal cortex | 10.96906286 | 11.50721682 | 2.272692303 | 13.9113121 | 0.800077045 | 0.434153297 | 1.816524389 | 1.141162804 |
| **Female** | mHFD | D2F5 | Prefrontal cortex | 10.53635025 | 9.40790616 | 2.170723838 | 12.07610391 | 0.128057381 | 0.823533695 | 1.800838875 | 0.569184786 |
| **Female** | mHFD | D4F6 | Prefrontal cortex | 9.845253707 | 11.34150377 | 1.948731391 | 12.12850232 | 0.100289774 | 0.493863618 | 2.703782334 | 0.37103511 |
| **Female** | mHFD | D6F6 | Prefrontal cortex | 9.640620714 | 7.679691318 | 1.877760896 | 8.438375257 | 0.390746718 | 1.52576645 | 2.964967327 | 0.74469653 |
| **Female** | mHFD | D8F5 | Prefrontal cortex | 12.32557532 | 11.79581484 | 2.489663434 | 8.944079507 | | | | |
